# VUSVista: Enhancing the Curation of Variants of Uncertain Significance

**DOI:** 10.64898/2025.12.02.691669

**Authors:** Esther Spiteri, Jeanesse Scerri, Jean-Paul Ebejer

## Abstract

**Background:** The increased demand for genomic sequencing has consequently led to a rise in the number of Variants of Uncertain Significance (VUS), which are classified as such due to insufficient evidence to determine their pathogenicity. VUS lead to incomplete diagnosis for patients, potentially causing anxiety due to the uncertainty associated with VUS and resulting in suboptimal management.

**Results:** We developed a software, VUSVista, for organised and semi-automated VUS curation. It offers American College of Medical Genetics and Genomics (ACMG) criteria management, external database references, standardised Human Phenotype Ontology (HPO) terms for sample phenotypes, links between variants and pseudonymised samples as well as audit trails. VUSVista automatically checks for updates to ClinVar’s germline classification entries of recorded VUS and searches for newly released publications referencing its VUS using LitVar 2.0. Users are notified whenever a ClinVar entry is updated or a relevant publication is found, helping them stay informed about new scientific discoveries. Assessing the significance of this new evidence may lead to VUS reclassification through VUSVista. One-to-one sessions and a focus group were conducted with medical laboratory scientists and a senior pharmacist to validate the system.

**Conclusions:** Overall, VUSVista facilitates VUS curation and reclassification, increasing the likelihood that patients receive a comprehensive diagnosis and the most appropriate treatment. Its implementation is publicly available on GitHub (https://github.com/estherspiteri/VUSVista).

## 1 Background

Clinical classification of genomic variants is crucial for determining their pathogenicity in relation to a disease or phenotype. The ACMG guidelines are commonly used in diagnostic labs to standardise the classification of variants in Mendelian disorders into five categories: “pathogenic”, “likely pathogenic”, “uncertain significance”, “likely benign”, and”benign”, based on variant evidence [1].

VUS result from a limited understanding of a variant’s contribution to a disease and their impact on protein function or phenotype expression [2]. Ackerman [3] refers to these variants as a *“genetic purgatory”* as they complicate disease diagnosis [4, 5]. Meanwhile, Giri et al. [6], Caswell-Jin et al. [7], and Tatineni et al. [8] observed that VUS are more prevalent among individuals of non-European descent, highlighting the healthcare inequalities faced by these populations. Notably, the adoption of Next-Generation Sequencing (NGS) technologies has led to a growing number of VUS awaiting reclassification, a task that is often delayed as laboratories prioritise sequencing new genetic samples for patients awaiting diagnosis. However, VUS reclassification is key to enhancing the impact of genetic testing on individuals [9]. VUS must be routinely re-evaluated to update their classification as new evidence emerges [4]. To address this need, our software, VUSVista, streamlines this process by automatically checking for new evidence and ensuring prompt re-evaluation through automated notifications.

Staying updated with the latest research is crucial to ensure variant pathogenicity classifications reflect current genomic discoveries. The rate of literature publication is steadily increasing due to advancements in sequencing technologies [10], creating a need for automatic text-mining procedures to manage this growing data [11, 12]. At least 79% of genomic variants mentioned in publications indexed by PubMed and PubMed Central Open Access Subset (PMC-OA) [13] are presented in a non-standardised form [14]. An example of the latter includes the most commonly used genetic variant form: “*wild type amino acid (w) + location number (N) + mutated type amino acid (m)*”, for example V600E, instead of standardised nomenclature such as the Human Genome Variation Society (HGVS) nomenclature [14]. Consequently, as shown in Figure 1, PubMed returns incomplete results, as it fails to recognise variant aliases and retrieves only exact matches [14]. Recognising entities, such as variant names, in text is known as Named-Entity Recognition (NER), while Named-Entity Normalisation (NEN) maps them to identifiers like the variant’s Reference SNP Identification (rsID) or HGVS form [14]. LitVar 2.0 is an actively maintained, publicly available semantic search engine which performs NER and NEN on its knowledge base (including PubMed, PMC-OA and articles’ supplementary material) as well as on the user’s query, enabling more comprehensive results [15, 16]. It is provided by National Center for Biotechnology Information (NCBI), thereby making it a reliable resource for use within VUSVista to retrieve publications referencing recorded VUS. Recent advances in Large Language Models (LLMs) make literature search and variant normalisation well suited to hybrid approaches that combine traditional methods, like those used by LitVar, with LLM-based interpretation. This combination opens up new possibilities for progress in genomic variant extraction and interpretation.

**Fig. 1.**
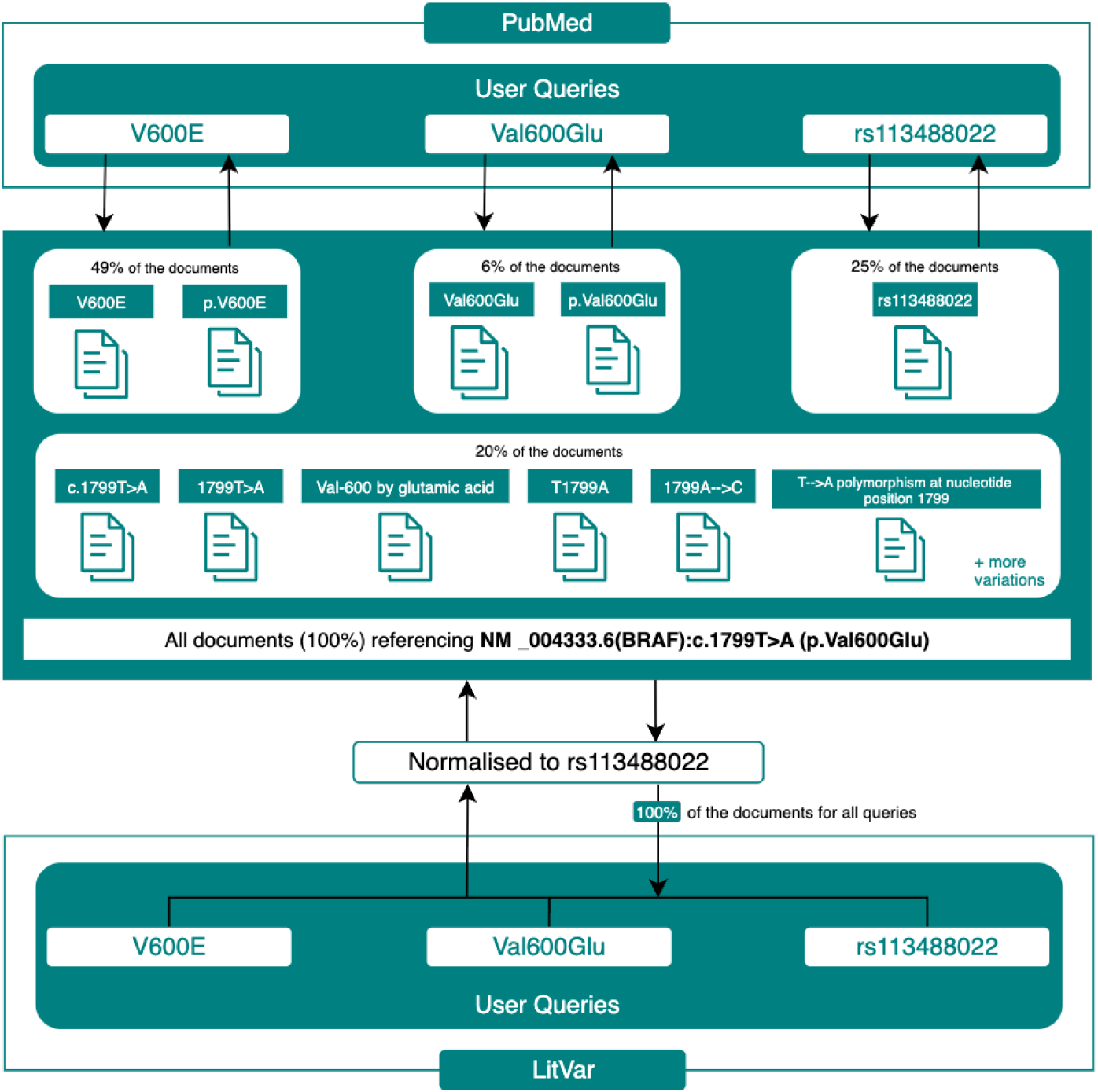
Implications of NER and NEN in literature search engines for genomic variant search. General search engines such as PubMed rely on exact or partial keyword matching, so multiple queries with different nomenclatures of the same variant are needed to find all relevant documents, which may lead to an incomplete literature search. In contrast, systems such as LitVar, which apply NER and NEN, enable comprehensive retrieval of all publications referencing a variant with a single query by normalising variant names in both the literature and the user’s query. Adapted from [14].

## 2 Implementation

VUSVista is a software tool for VUS curation that automatically monitors scientific updates to aid reclassification. To validate the system, one-to-one sessions and a focus group were conducted with medical laboratory scientists and a senior pharmacist as explained in Supplementary Section 4 in accordance to the University of Malta’s URECA Research Ethics and Data Protection (REDP) system (application ID: CMMB-2023-00010). The project’s implementation is available on GitHub.^1^ Additionally, a user manual for the software can be found in the Supplementary Information.

### 2.1 VUSVista Architecture

VUSVista is composed of three main entities: the PostgreSQL database, the Python server and the React frontend, as shown in Figure 2. The database stores all VUS-related information associated with the GRCh37 build, ACMG criteria, publication data and more, whilst the Python server functions as the Application Programming Interface (API) for VUSVista’s logical operations. VUSVista can be adapted to GRCh38 by obtaining the corresponding GTF file and updating the build from the configuration file. The frontend’s React application consists of different user-acessible views which registered users can navigate to after logging in.

**Fig. 2.**
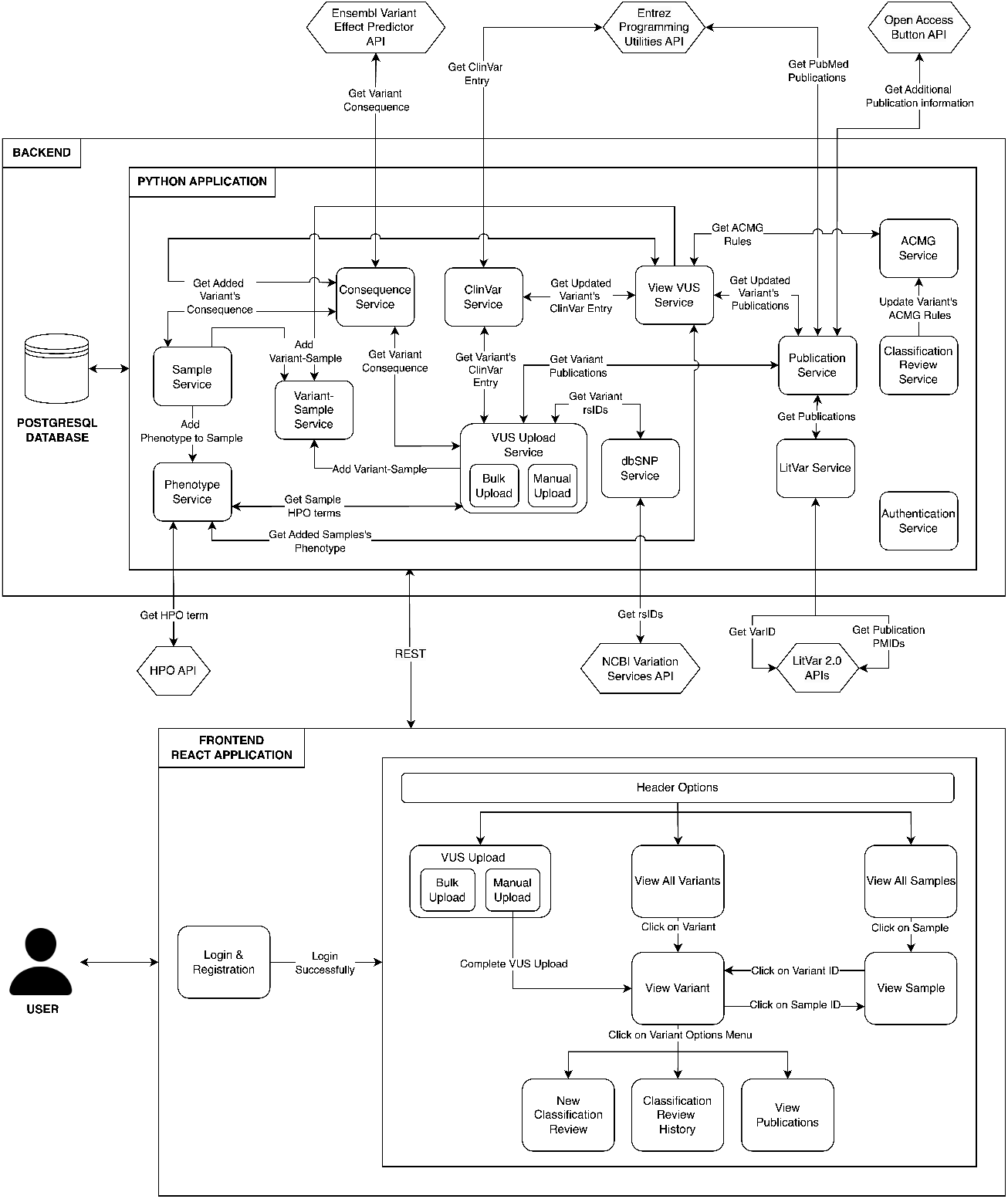
A high-level system architecture displaying the backend and frontend entities that make up VUSVista. The backend is composed of a PostgreSQL database and a Python application which contains services that communicate with one another. Hexagonal shapes represent external APIs that are publicly accessible whilst the term “Variant-Sample” refers to the occurrence of a variant in a sample. The backend and the frontend interact through Representational State Transfer (REST) calls. The frontend is a React application that serves as an interface for VUSVista’s users. It comprises user-accessible views that allow users to make use of the system’s features. The “Header Options” are the navigation options accessible from VUSVista’s website header that are visible at all times to authorised users.

### 2.2 Database Schema

Figure 3 displays VUSVista’s database schema through an Entity Relationship Diagram (ERD). This diagram represents the database’s structure indicating the declared entities and their respective attributes. Relationships between these entities are also displayed with their types clearly marked and explained in the diagram’s legend.

**Fig. 3.**
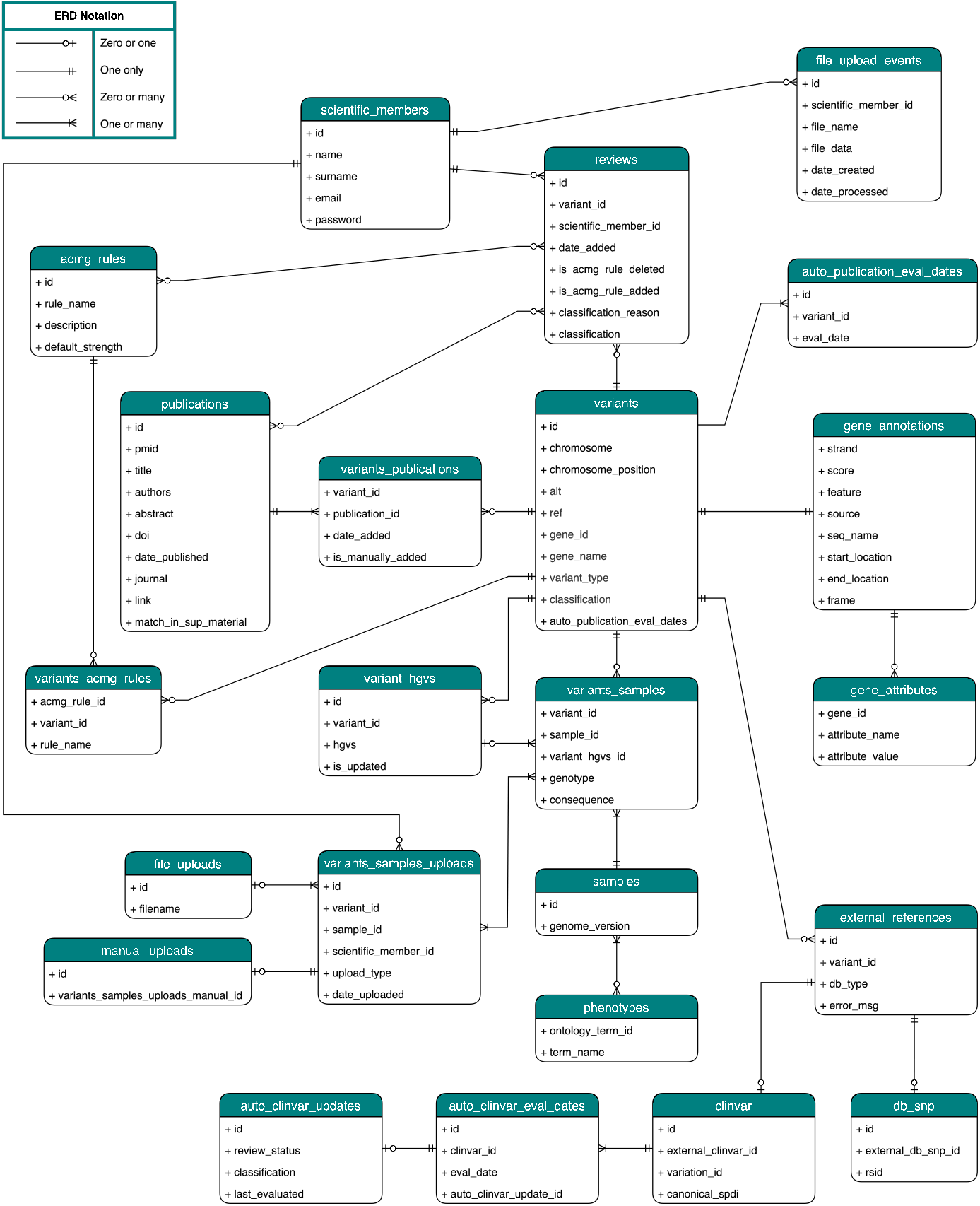
Entity Relationship Diagram (ERD) for VUSVista’s database.

### 2.3 User Registration and Login

In order to ensure that VUSVista is accessed only by qualified scientific personnel in a clinical or diagnostic settings with access to genomic data, user registration is exclusively managed by the system administrator. Passwords are salted and then hashed using the SHA-256 hash function prior to being saved for secure storage in VUSVista’s database. The Python library Flask-Login (version 0.6.2) is then used to manage user sessions.

### 2.4 VUS Submission

One of the most critical features of VUSVista, is its variant submission functionality. Users can carry out bulk uploads of variants or only submit a single variant. In order to record a variant in VUSVista, users are required to provide the information shown in Supplementary Table S1, based on the GRCh37 genome build, for both submission types. The system uses the HPO [17] to describe patient phenotypes, ensuring standardised nomenclature throughout.

#### 2.4.1 Bulk Upload View

The bulk upload submission method is most beneficial when initially setting up VUSVista and populating it with all the VUS entries that the laboratory or clinic has already recorded. Bulk upload requires users to submit an Excel file with specific column names and data formatting rules in the bulk upload view which is displayed in Supplementary Figure S1. A file template, shown in Figure 4, is provided for system users to use as a guide.

**Fig. 4.**
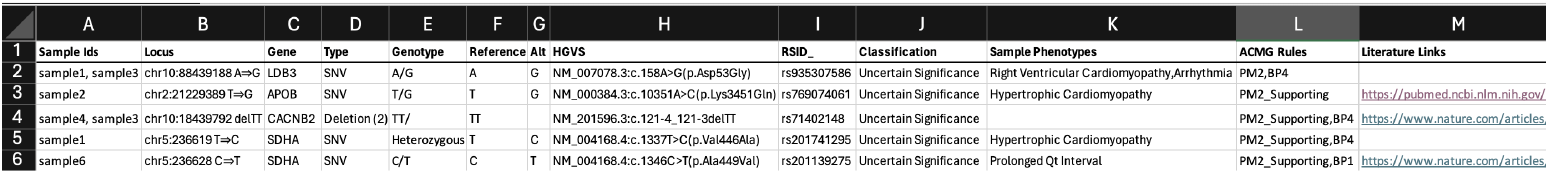
VUSVista’s Excel file upload template.

File processing takes place asynchronously through a file upload scheduler which runs a background job periodically without interrupting the main application’s execution. The frontend then polls the backend to check if this task is complete.

#### 2.4.2 Manual VUS Submission View

To address the need for users to add individual VUS, a manual VUS submission functionality has been implemented. An input form, displayed in Figures 5 and 6, guides the user to provide all mandatory variant data and optional information through its intuitive design. Prior to submission, VUSVista displays a summary of the variant the user wishes to store in its database as a verification step.

**Fig. 5.**
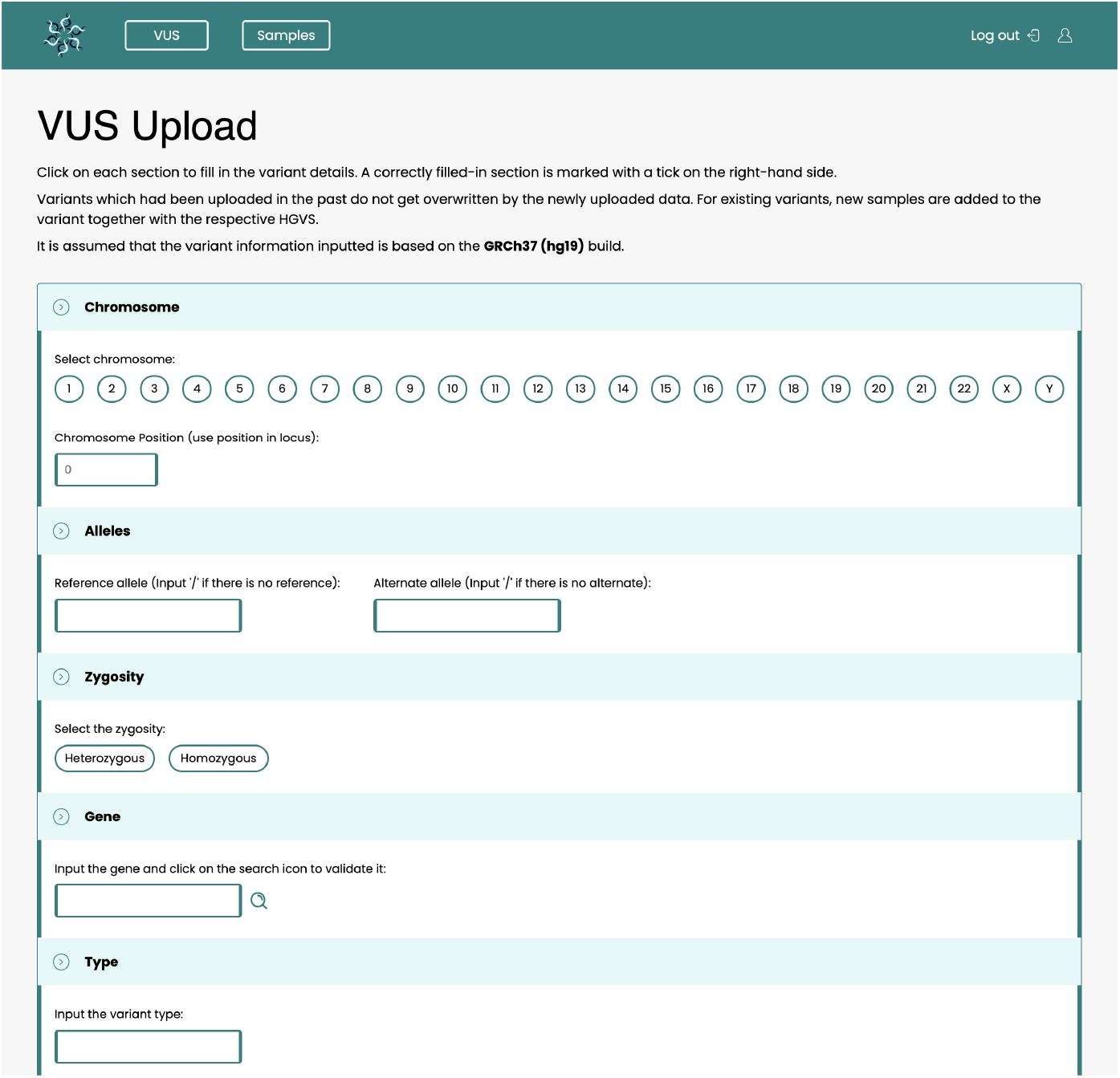
Screenshot from VUSVista of manual submission form (part 1). All fields visible in this figure are mandatory. Reference and alternate alleles can be set to “/” in the case of insertions or deletions. Gene symbols need to verified after input. VUSVista ensures that these genes are found in the GRCh37 annotation file. Examples of variant types are: “SNV”, “insertion”, “deletion”.

**Fig. 6.**
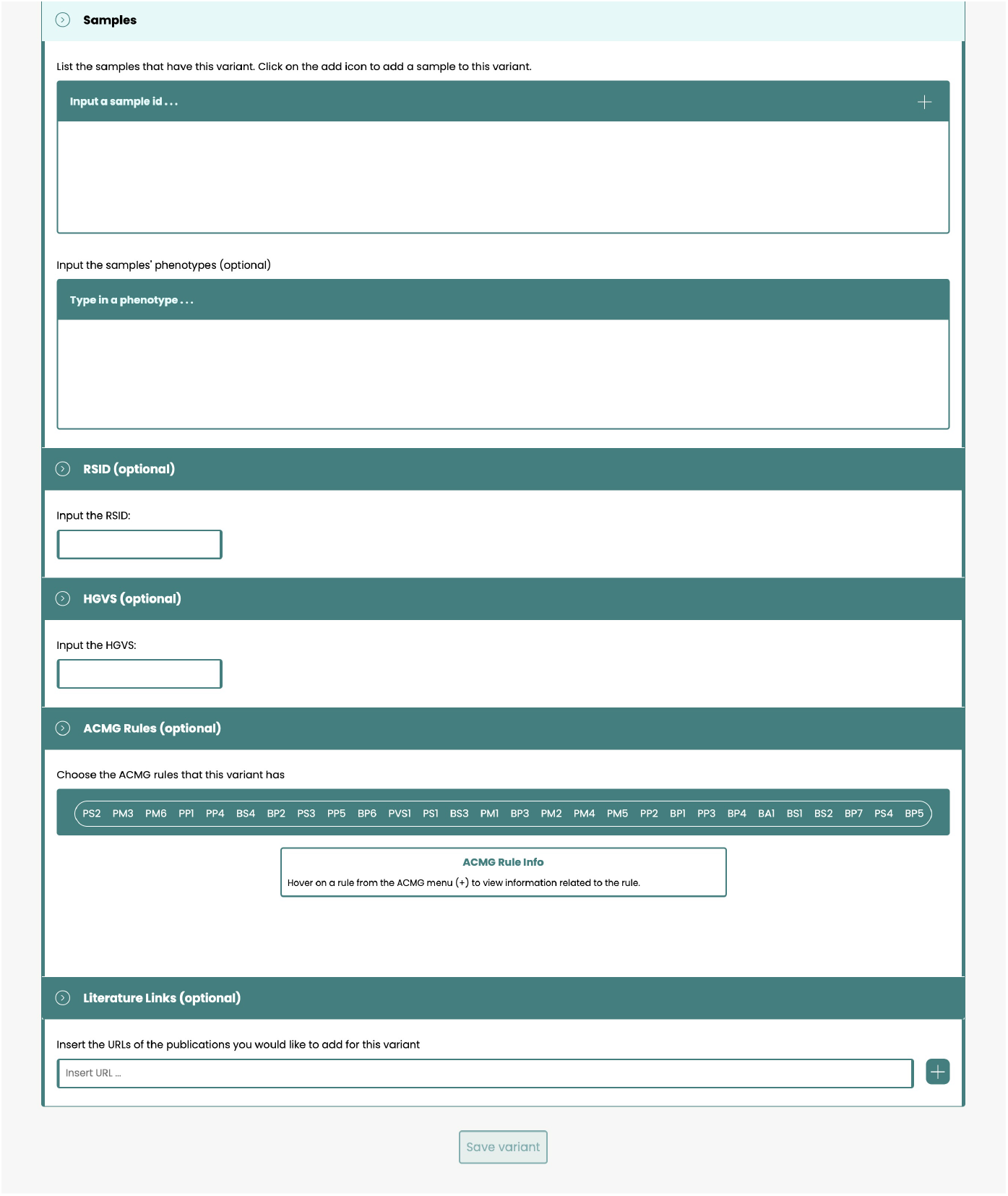
Screenshot from VUSVista of manual submission form (part 2). Users are required to input at least one sample identifier. The rest of the fields are optional. When typing a phenotype, a dropdown with HPO term suggestions is shown to the user from which they can select from. Hovering on an ACMG criteria, displays information explaining what it represents and its strength.

#### 2.4.3 Variant Processing

The variant processing procedure is displayed in Supplementary Figures S2a and S2b. Initially, the submitted variants’ external references are retrieved from Database for Single Nucleotide Polymorphisms (dbSNP) [18] and ClinVar [19]. Next, the system fetches the consequences of each variant, that is their impact on proteins, transcripts, and regulatory elements such as those affecting gene expression. This is followed by the conversion of sample phenotypes to HPO terms. For variants already recorded in VUSVista, the provided samples are merged with the existing ones. Otherwise, samples for new variants are stored in the system’s database. The newly discovered variants themselves and their external references are then also recorded in the database. Publications that reference the variants, that are either retrieved from LitVar 2.0 or provided by the user, are then handled by the system. These publications and the variants’ ACMG criteria are then stored in VUSVista’s database. The relationship between every variant and its samples is then recorded in the database with information that is sample-specific such as the genotype, the HGVS and the variant’s consequence.

### 2.5 Variants

The primary purpose of VUSVista is to record and manage VUS. Users can view, filter, and update variants, along with related publications, samples carrying these variants, and phenotypes observed in the patients of those samples. The system also attempts to link variants to dbSNP and ClinVar and it regularly checks for ClinVar updates or newly released publications referencing the variant. Together, these features provide VUSVista users with deeper insights into VUS.

All variants recorded in VUSVista are initially submitted as VUS. However, some may eventually be reclassified and no longer hold this classification.

### 2.5.1 Variant View

The variant view first displays a summary of the variant’s information as shown in Figure 7.

**Fig. 7.**
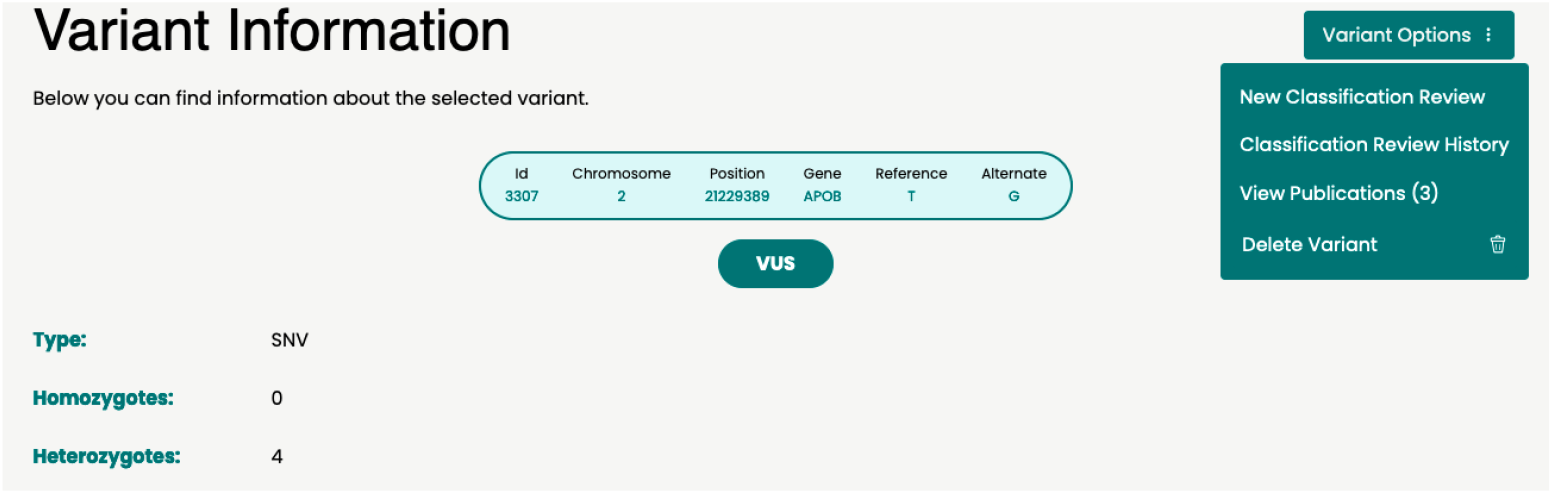
Screenshot from VUSVista of variant view’s variant details summary.

This is followed by an external references area which contains variant information related to dbSNP and ClinVar. Both the rsID and the ClinVar entry can be either confirmed as the corresponding entry for the variant (Figure 8) or suggested as a possible match by VUSVista (Figure 9). The latter occurs whenever there is a mismatch between the external reference’s variant and the user-inputted variant. The rsID can be manually updated. In this scenario, VUSVista removes any automatically added variant publications since they are related to the old rsID. It then looks up publications and attempts to find a ClinVar entry, both in relation to the new rsID.

**Fig. 8.**
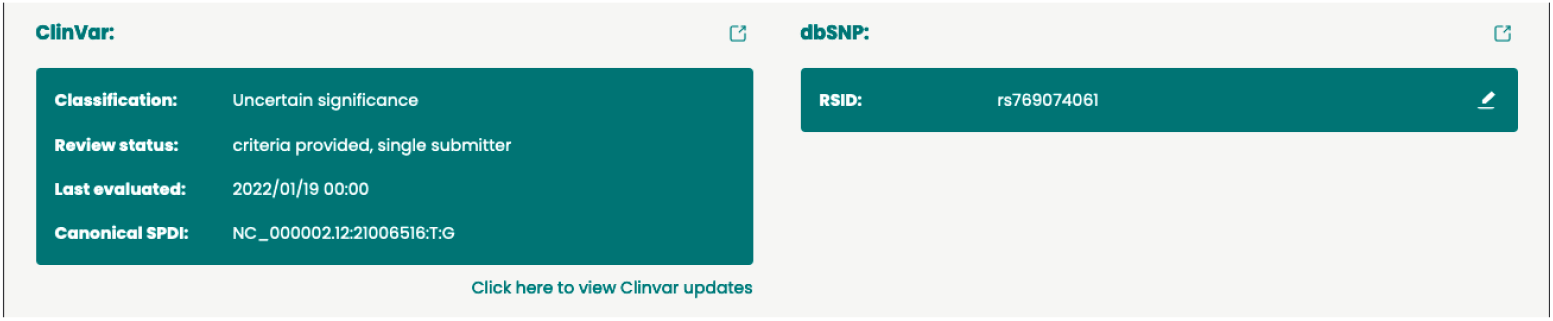
Screenshot from VUSVista of valid variant external references. VUSVista’s variant information matches that of dbSNP and ClinVar.

**Fig. 9.**
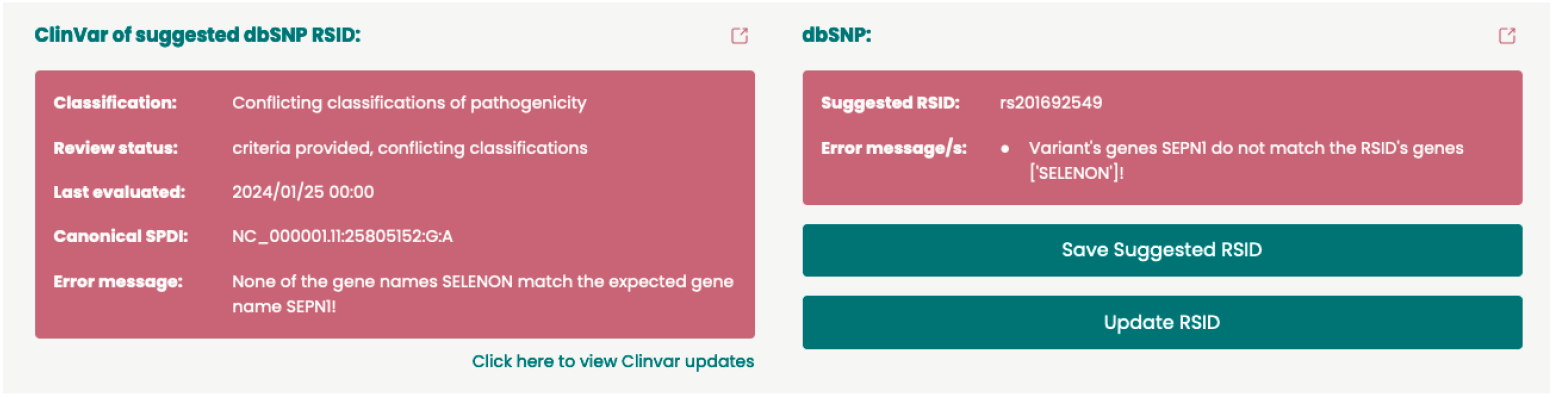
Screenshot from VUSVista of suggested variant external references. VUSVista’s variant information does not exactly match that of dbSNP and ClinVar.

One of the software’s most beneficial features for variant reclassification is its ability to automatically check for any clinical classification updates for every variant which has a ClinVar entry. In the external references area, an audit trail is provided through a calendar displaying marked dates of when these automatic checks are performed, along with a timeline indicating when and what changes actually occurred. This audit trail is shown in Supplementary Figure S3.

VUSVista users can add or remove an ACMG criterion from a variant through the variant view as demonstrated in Figure 10. In both cases, the user is required to fill out a classification review form which is prepopulated with the respective ACMG criterion. In this form, the user can also revise the variant’s classification whenever the updated ACMG criterion changes the existing classification, in line with ACMG guidelines. It enables users to keep track of criteria changes that are made to the variant since first being recorded in VUSVista.

**Fig. 10.**
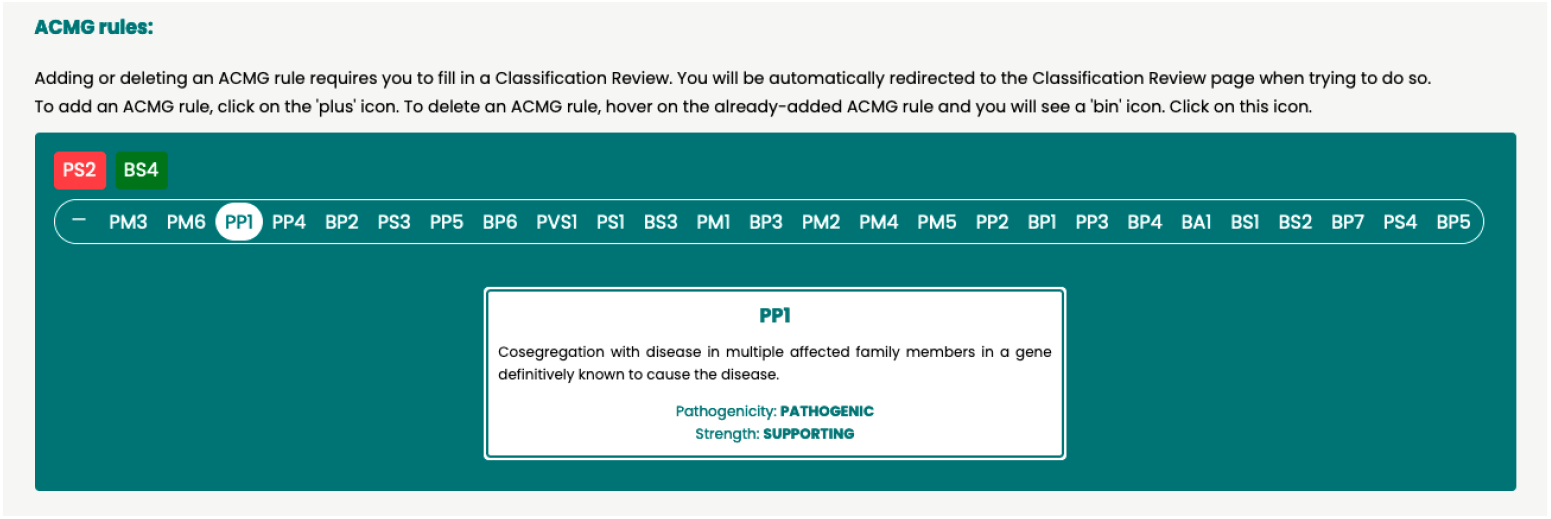
Screenshot from VUSVista of the variant view’s ACMG criteria area. Upon hovering over an ACMG criterion, its strength and an explanatory description is displayed. Users can update the variant’s ACMG criteria from this area. Both adding and removal of ACMG criteria require the user to submit a classification review to record the action.

Samples exhibiting a variant are displayed in a searchable table, shown in Figure 11. For each sample, the HGVS nomenclature representing the transcript on which the variant is located is shown, if provided. The variant consequence, which is based on the transcript, is also visible.

**Fig. 11.**
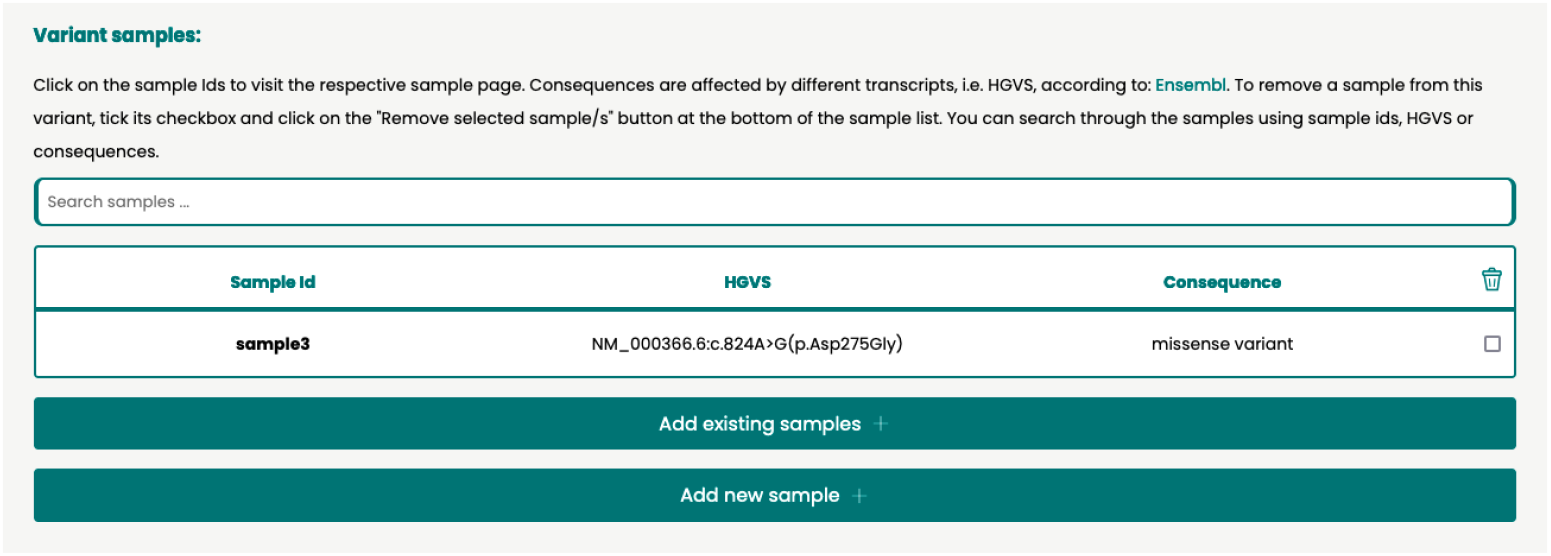
Screenshot from VUSVista of the sample area in the variant view. It lists all samples exhibiting the variant together with HGVS nomenclature, if provided by the user, and the variant’s consequence (based on the HGVS).

The functionality to remove a sample from a variant is also available, requiring user confirmation. A variant remains stored in VUSVista’s database even if all of its samples are removed, unless it is deleted by a user. Additionally, users can choose to add either previously recorded samples from VUSVista or new ones to the variant’s sample list.

The variant view also contains a list of all the unique HPO terms that describe the phenotypes exhibited by the variant’s samples (Figure 12). Users can access the HPO‘s website when clicking on a phenotype and they can also run a search for publications that mention both the variant and one of the phenotypes.

**Fig. 12.**
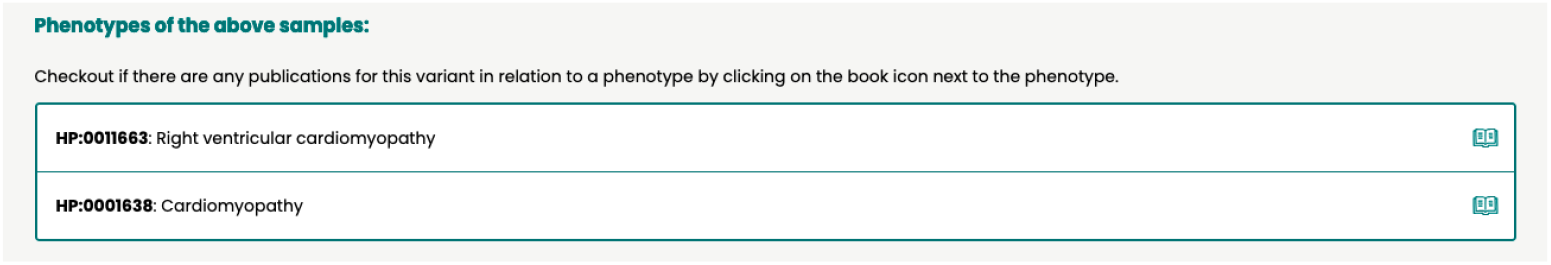
Screenshot from VUSVista of all samples’ phenotypes in the variant view.

When a user decides to delete a variant, VUSVista prompts them to confirm their decision to prevent accidental deletions. Variant deletion results in the removal of any information related to it.

#### 2.5.2 Classification Review

The purpose of a classification review is to allow users to change a variant’s classification, for example from VUS to “likely pathogenic”, provided that supporting evidence is available. Another use case is to record some evidence, that is deemed to be informative by VUSVista users, that might contribute to a change in the variant’s classification in the future.

##### Classification Review View

In order to submit a classification review, users must fill out the classification review form shown in Figure 13. This form requires users to populate one or more of the following as supporting evidence:

**Fig. 13.**
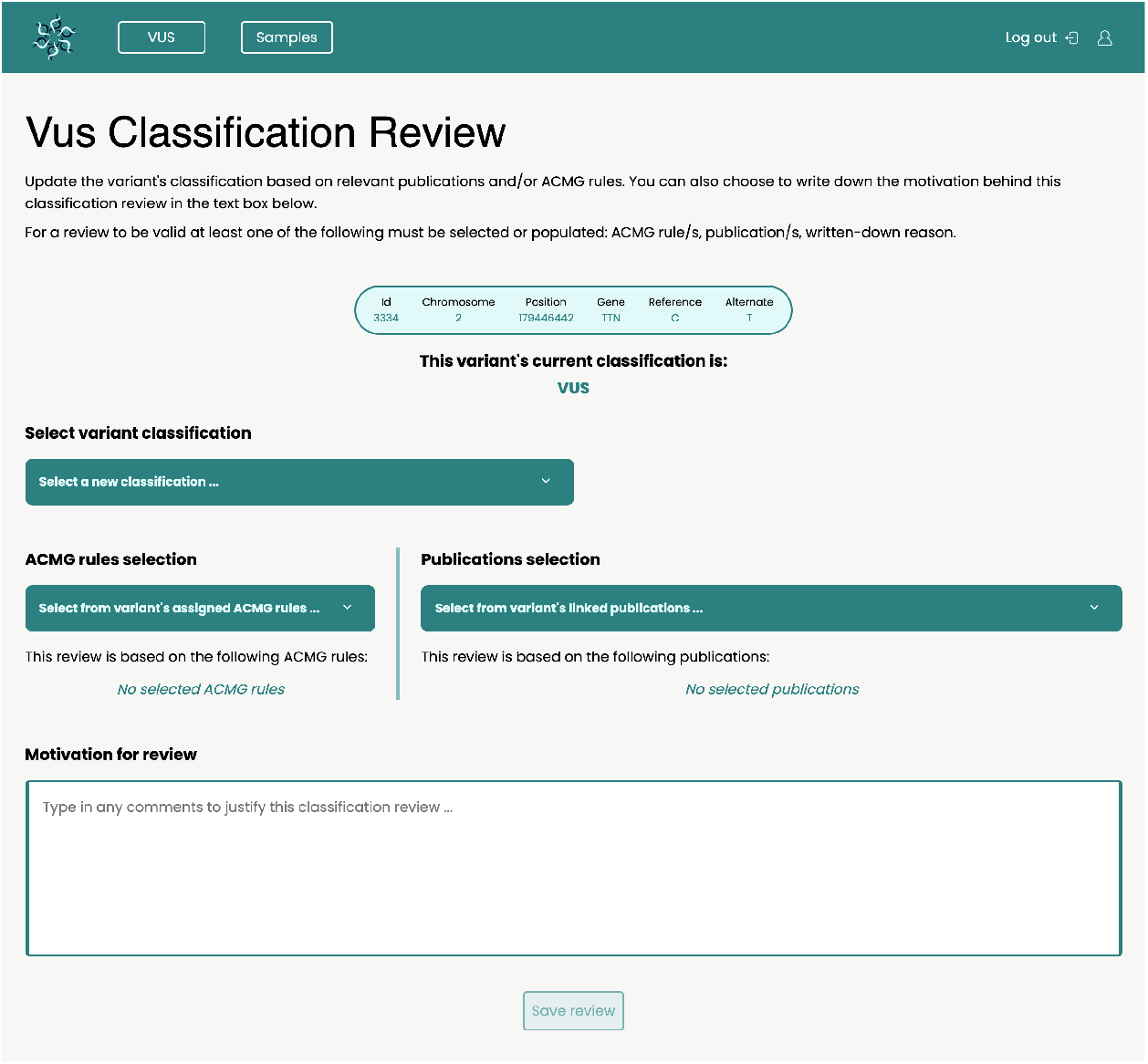
Screenshot from VUSVista of an example classification review form.

- ACMG criteria that are already assigned to the variant;
- publications recorded in VUSVista that reference the variant; or
- a free-text field for explaining the rationale behind the review submission.

##### Classification Review History View

Any classification review that is created, is then recorded in the classification review history, as shown in Figure 14. This history displays an audit trail timeline starting from when the variant was first submitted to VUSVista to the latest submittted classification review form. Such audit trails contribute to the Quality Management System (QMS) of the organization adopting VUSVista by providing a structured record of variant management activities whilst promoting transparency, traceability and communication amongst staff.

**Fig. 14.**
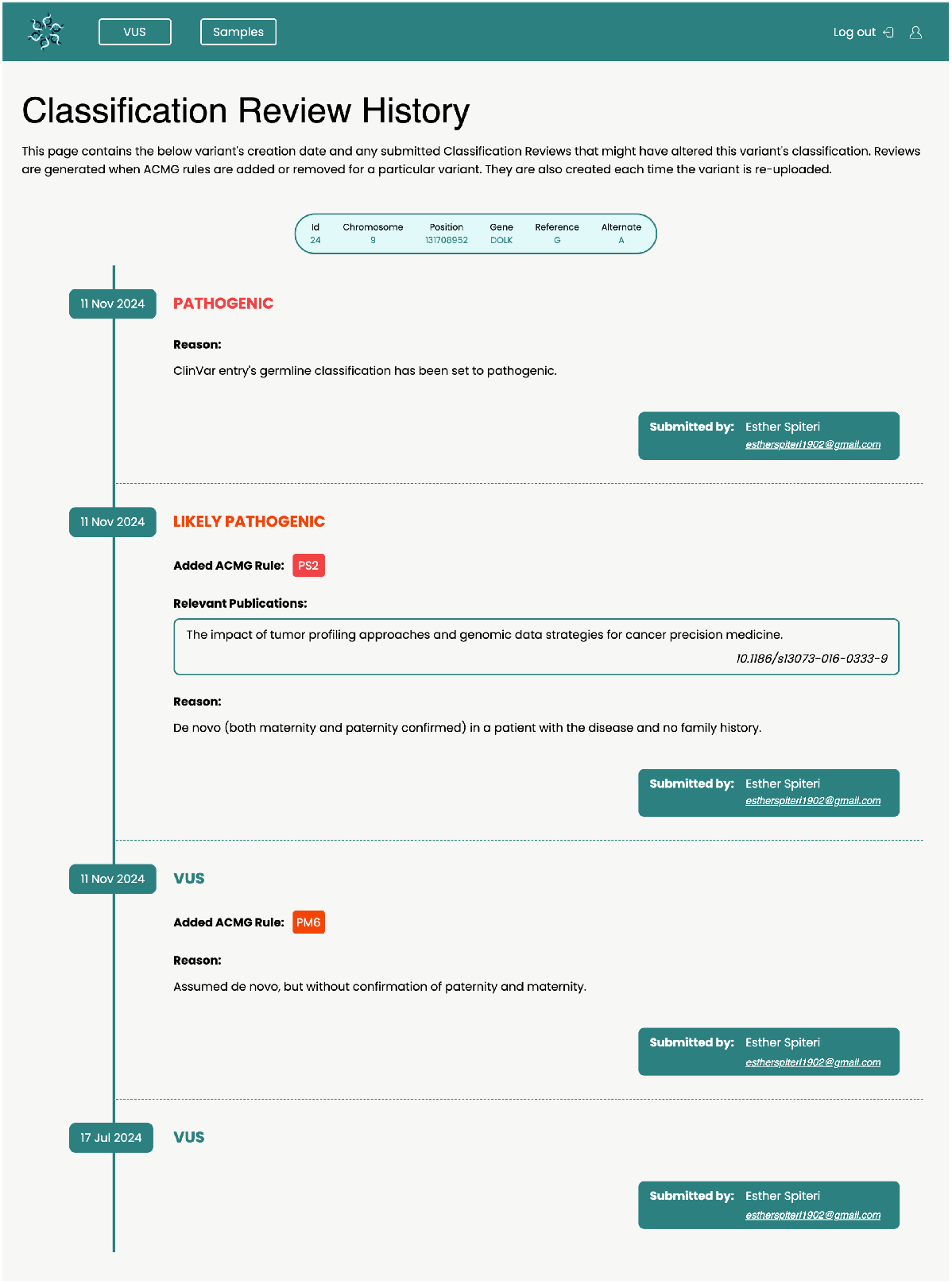
Screenshot from VUSVista of an example of a classification review history, using hypothetical data where the cited publications may not necessarily impact the variant’s classification.

#### 2.5.3 Publication View

Each variant in VUSVista has a publication view listing literature referencing the VUS, including publications automatically found via LitVar 2.0’s API and those manually added by users. Ideally, these publications have a Digital Object Identifier (DOI) as it is used to identify each publication in VUSVista and eliminate duplicates. Publication information then indicates whether the variant is mentioned within the publication’s supplementary material or in its main text, as is the case in Figure 15.

**Fig. 15.**
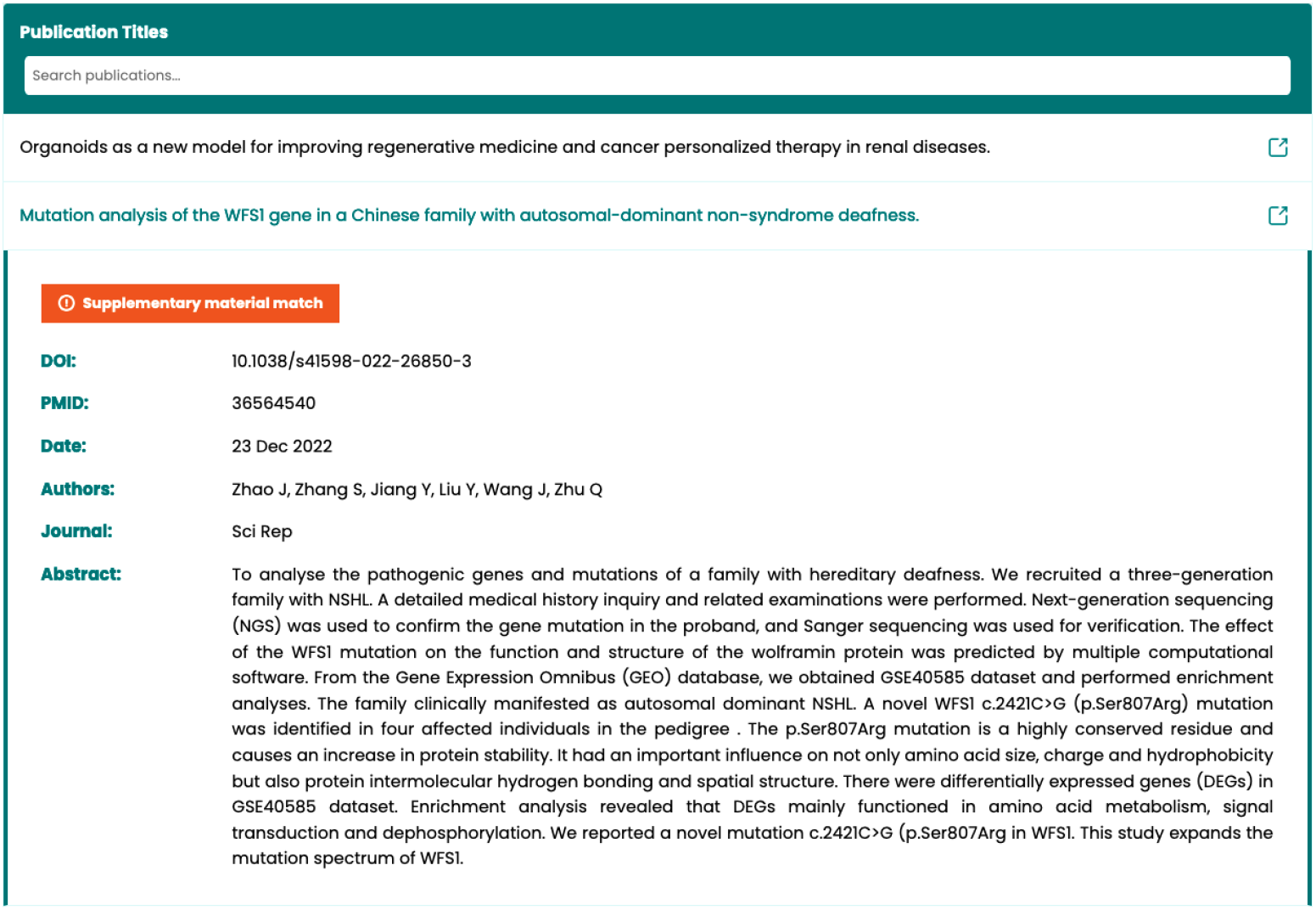
Screenshot from VUSVista of information for a publication which references the variant in its supplementary material.

Furthermore, from the publication view, one can generate a report that includes all of the variant’s publications along with any retrieved information about them. This report can be shared with clinicians, scientific members of staff, or external diagnostic laboratories to help expand their knowledge on a particular VUS they may have encountered.

A critical feature of VUSVista is that it can automatically check for any newly released publications that reference variants stored in VUSVista using LitVar 2.0. An archive view, serving as an audit trail, is accessible from the publication view and is shown in Figure 16. It contains a calendar showing when these automatic checks are performed, along with a timeline indicating when and which publications were added.

**Fig. 16.**
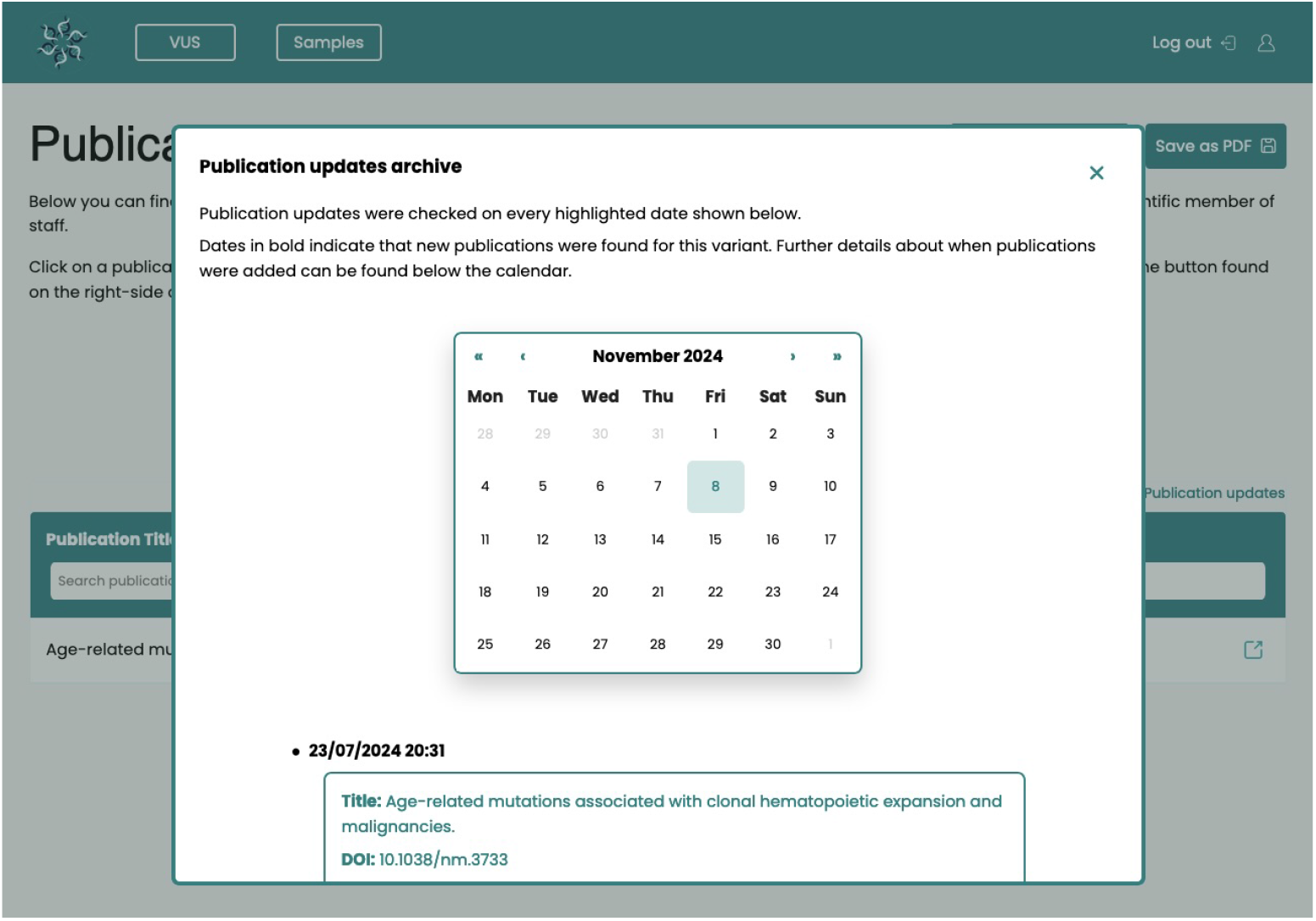
Screenshot from VUSVista of the publication updates archive consisting of calendar marking automatic checks for newly released publications, that reference the variant, as well as a timeline showing what publications were found.

### 2.6 Samples

Information regarding samples that have at least one variant stored in VUSVista are mostly vital for when a VUS’ classification changes to “likely pathogenic” or “pathogenic”. This is because authorised users can trace back to whom the variant’s samples belong to, by accessing the internal laboratory records, and providing guidance to genetic counsellors and consultants, who can then consult with the affected individuals and their families.

#### 2.6.1 Sample View

The sample view contains all the information associated to a single sample. From this view, users can add new HPO terms or remove existing ones from the sample. The selected HPO terms will help VUSVista users better understand the potential impact a VUS might have on the affected patient from whom the sample was taken.

All the variants exhibited by the sample are shown in the variant area (Figure 17). These can be filtered by any visible variant information, including the HGVS nomenclature. The HGVS field is editable to correct input errors, and any correction is automatically applied to all of the sample entries that share the same HGVS value.

**Fig. 17.**
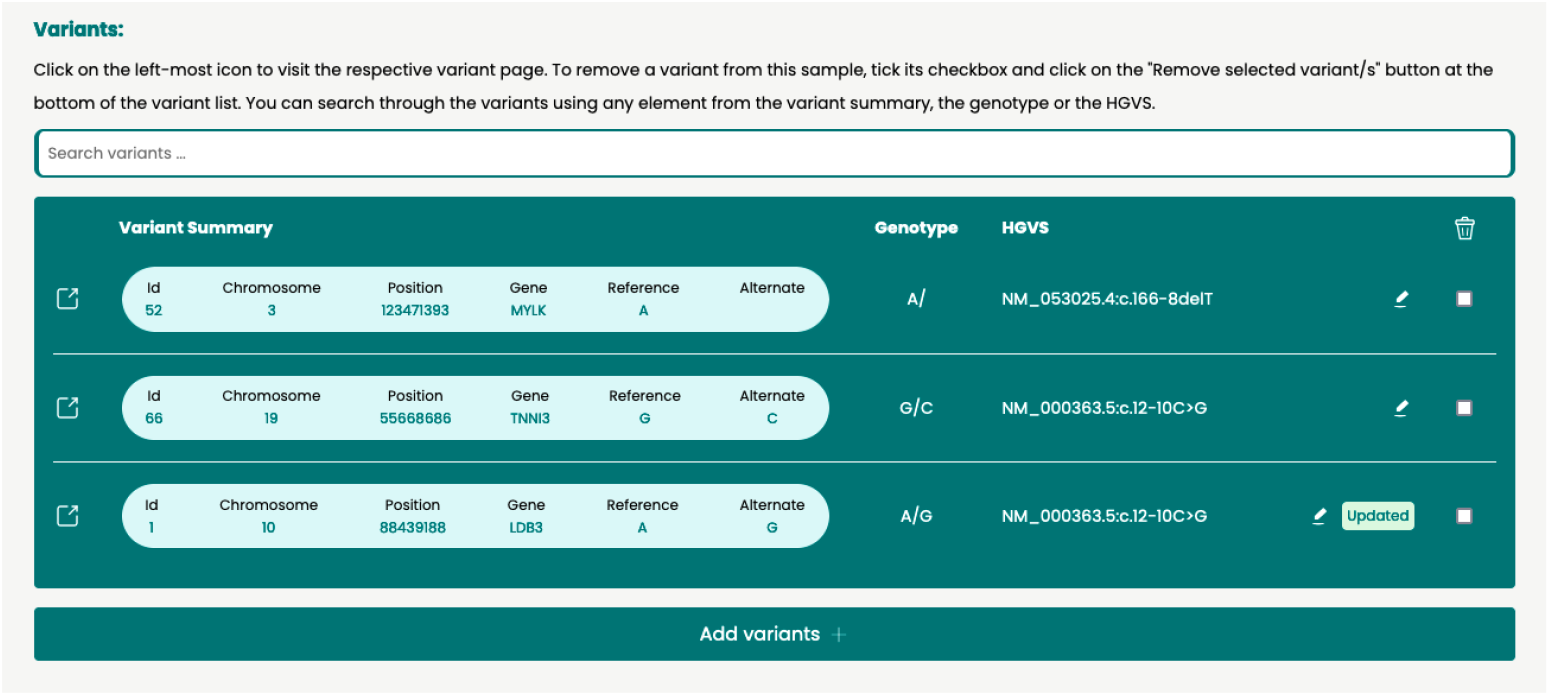
Screenshot from VUSVista of variants exhibited by sample in sample view. The variant view can be accessed by clicking on the leftmost icon found in every variant entry. The HGVS can be edited by clicking on the “pen” icon.

If the sample starts exhibiting a VUS that has already been submitted to VUSVista, users can “add” it to the sample. Variants can also be removed from a sample after confirmation. If all of the sample’s variants are removed, then the sample is automatically deleted from VUSVista together with all of its information. Samples may also be deleted from their respective sample view. Variants that end up having no samples are not removed from VUSVista unless they are deleted by the user. This ensures that no vital variant information is removed without the user’s consent and supports the preservation of the variant’s audit trail, which is especially useful when new samples exhibit the same variant.

### 2.7 Automated Checks

The following checks are done automatically by VUSVista at regular set intervals. Whenever new findings are discovered, all of the system’s registered users receive an email notifying them of the update. This notification feature allows them to act on new evidence related to VUS that might potentially result in variant reclassifications. Moreover, VUSVista’s automated checks for new discoveries from the scientific community save researchers time by removing the need to manually search for updates, thereby reducing their workload.

#### 2.7.1 ClinVar

By default, VUSVista retrieves all the recorded variants with a ClinVar entry from the system once a week and checks for updates on the following:

- the last date on which the germline classification is evaluated
- the germline classification itself and
- the review status of the submitted classification (the assertion criteria)

The process of checking for ClinVar entry updates is demonstrated in Supplementary Figure S4 and can be configured to run at a different interval. If one or more variants’ ClinVar entries are updated, then a notification email, shown in Supplementary Figure S5, is sent to all registered users.

#### 2.7.2 Publications

VUSVista utilises LitVar 2.0 to automatically search for newly released publications every ten days by default, as illustrated in Supplementary Figure S6, with this interval easily configurable. This search is done for every variant with a valid rsID or, alternatively, with one of its HGVS nomenclature. PubMed is then queried in order to gather information about the returned publications. Newly discovered publications for a variant are merged with the variant’s already recorded publications based on their DOI. If new publications are found for at least one variant, an email is sent out to all registered users, as shown in Supplementary Figure S7.

## 3 Results and Discussion

In this section, we present the features that make VUSVista a valuable tool for individuals who encounter VUS in their daily work. We also go over the system’s limitations and potential areas for improvements to provide a transparent and comprehensive review.

### 3.1 VUSVista’s Notable Features

During the focus group conducted as part of the system’s evaluation, participants praised VUSVista’s interface as intuitive, especially for users without an IT background. Moreover, they described it as self-explanatory and easy to manage.

A significant feature of VUSVista is its ability to perform regular checks of both ClinVar and newly released publications for updates related to the stored VUS. When an update is detected, the system automatically sends an email notification to all of its registered users. This functionality is particulary beneficial for variant reclassification since it keeps VUSVista users informed about the latest findings without the need to carry out manual searches, thereby reducing their workload.

Another notable feature of VUSVista is its ability to link variants with the samples that exhibit them. This allows users to promptly notify patients associated with samples affected by variant classification updates. As a result, patients’ treatments can be adjusted as necessary without requiring VUSVista users to consult intermediate databases to link variants to samples.

Additionally, VUSVista’s pseudonymisation aspect is crucial for GDPR compliance, as it ensures that only authorised laboratory personnel with access to a patient-sample directory can identify patients. This feature is of utmost importance to protect the patients’ privacy so that no sensitive data is linked to any individual by unauthorised VUSVista users. Thus, our system is in line with ethical standards and data protection rules.

### 3.2 Identified Gaps and Areas for Improvement

VUSVista queries ClinVar by rsID, so it does not distinguish variants at the same locus since an rsID identifies a locus and not a unique variant. The system could allow users to update linked ClinVar entries and notify them when multiple ClinVar records share a single rsID. It would then need to filter out those papers mentioning the selected variant using machine-learning techniques.

Notably, the system’s variant audit trail in the classification review history does not record sample changes which should be logged for traceability and accountability.

VUSVista’s VUS submission process could be improved by supporting ACMG criteria with intermediate strengths, such as PM2 Moderate, and by providing translations between the GRCh37 and GRCh38 builds. The system could also record the gender for each sample to enable gender based filtering, for example in infertility studies, and flag proband samples to support segregation analysis with phenotype data. These additions would enhance variant interpretation while maintaining patient privacy.

Finally, VUSVista does not query ClinVar again for variants that lacked a ClinVar entry when initially recorded. Adding scheduled checks would improve completeness.

## 4 Conclusions

In a diagnostics laboratory or a clinical setting, maintaining a standardised VUS management system for ease of retrieval, continuous documentation and auditing is essential. In addition, scientific members of staff have the tedious responsibility of regularly checking if there is sufficient evidence available to reclassify any VUS.

We developed VUSVista, a VUS curation system that focuses on storing VUS in a standardised manner together with any other relevant information. Our system provides users with an audit trail of updates to a variant’s ACMG criteria alongside any of its classification changes, providing transparency and integrity to the stored data. For every recorded variant, VUSVista attempts to find an rsID from dbSNP and a corresponding ClinVar entry, as well as relevant publications referencing the respective variant. Moreover, VUSVista carries out automated checks periodically for updates in ClinVar’s germline classifications and for any newly released publications. It sends email notifications when an update is found, helping users stay informed and potentially update variant classifications based on the provided evidence. The afore-mentioned features have already attracted interest from local diagnostic molecular genetics laboratories, and we are confident that they will also appeal to international laboratories, particularly those looking for accessible solutions that reduce the need for extensive manual curation.

While VUSVista still has room for improvement and potential for future development, it can aid anyone who works with VUS, including scientific members of staff, clinicians and researchers, in their day-to-day work and potentially encourage VUS reclassification. In the end, this work aims to provide a more comprehensive diagnosis, ultimately improving the health and well-being of patients.

## Supporting information

VUSVistaManual.pdf

## Declarations

The authors declare **no competing financial interest**.

### Ethics approval and consent to participate

This study was approved by the URECA committee at the University of Malta in accordance with the university’s code of practice and research ethics review procedures. A Research Ethics and Data Protection (REDP) form was submitted for this purpose. (REDP Application ID: CMMB-2023-00010)

## 5 List of Abbreviations

ACMG: American College of Medical Genetics and Genomics
API: Application Programming Interface
dbSNP: Database for Single Nucleotide Polymorphisms
DOI: Digital Object Identifier
ERD: Entity Relationship Diagram
HGVS: Human Genome Variation Society
HPO: Human Phenotype Ontology
LLMs: Large Language Models
NCBI: National Center for Biotechnology Information
NEN: Named-Entity Normalisation
NER: Named-Entity Recognition
NGS: Next-Generation Sequencing
PMC-OA: PubMed Central Open Access Subset
REST: Representational State Transfer
rsID: Reference SNP Identification
QMS: Quality Management System
VUS: Variants of Uncertain Significance

## Supplementary information

This article has two accompanying supplementary files. One of which contains supplementary figures and tables (supplementary.pdf) whilst the other contains VUSVista’s user manual (VUSVistaManual.pdf).

## Acknowledgements

The authors would like to acknowledge the staff of the Molecular Pathology and Genetics laboratory, Pathology Department, Mater Dei Hospital, for their contribution to the individual and collective evaluation of this solution.

## Footnotes

1 https://github.com/estherspiteri/VUSVista

## References

[1] Richards S, Aziz N, Bale S, Bick D, Das S, Gastier-Foster J, et al. Standards and Guidelines for the Interpretation of Sequence Variants: A Joint Consensus Recommendation of the American College of Medical Genetics and Genomics and the Association for Molecular Pathology. Genetics in medicine : official journal of the American College of Medical Genetics. 2015;17(5):405–424.

[2] Kwong A, Ho CYS, Shin VY, Au CH, Chan TL, Ma ESK. How does reclassification of variants of unknown significance (VUS) impact the management of patients at risk for hereditary breast cancer? BMC Medical Genomics. 2022;15:122.

[3] Ackerman MJ. Genetic purgatory and the cardiac channelopathies: Exposing the variants of uncertain/unknown significance issue. Heart Rhythm. 2015;12(11):2325–2331.

[4] Bennett JS, Bernhardt M, McBride KL, Reshmi SC, Zmuda E, Kertesz NJ, et al. Reclassification of Variants of Uncertain Significance in Children with Inherited Arrhythmia Syndromes is Predicted by Clinical Factors. Pediatric Cardiology. 2019;40(8):1679–1687.

[5] Wan A, Place E, Pierce EA, Comander J. Characterizing variants of unknown significance in rhodopsin: A functional genomics approach. Human Mutation. 2019;40(8):1127–1144.

[6] Giri VN, Hartman R, Pritzlaff M, Horton C, Keith SW. Germline Variant Spectrum Among African American Men Undergoing Prostate Cancer Germline Testing: Need for Equity in Genetic Testing. JCO Precision Oncology. 2022;(6):e2200234.

[7] Caswell-Jin JL, Gupta T, Hall E, Petrovchich IM, Mills MA, Kingham KE, et al. Racial/ethnic differences in multiple-gene sequencing results for hereditary cancer risk. Genetics in Medicine: Official Journal of the American College of Medical Genetics. 2018;20(2):234–239.

[8] Tatineni S, Tarockoff M, Abdallah N, Purrington KS, Assad H, Reagle R, et al. Racial and ethnic variation in multigene panel testing in a cohort of BRCA1/2-negative individuals who had genetic testing in a large urban comprehensive cancer center. Cancer Medicine. 2022;11(6):1465–1473.

[9] Walsh N, Cooper A, Dockery A, O’Byrne JJ. Variant reclassification and clinical implications. Journal of Medical Genetics. 2024;p. jmg–2023–109488.

[10] Bugnon LA, Yones C, Raad J, Gerard M, Rubiolo M, Merino G, et al. DL4papers: a deep learning approach for the automatic interpretation of scientific articles. Bioinformatics. 2020;36(11):3499–3506.

[11] Leaman R, Wei CH, Allot A, Lu Z. Ten tips for a text-mining-ready article: How to improve automated discoverability and interpretability. PLOS Biology. 2020;18(6):e3000716.

[12] Singhal A, Simmons M, Lu Z. Text Mining Genotype-Phenotype Relationships from Biomedical Literature for Database Curation and Precision Medicine. PLoS computational biology. 2016;12(11):e1005017.

[13] Bethesda (MD): National Library of Medicine.: PMC Open Access Subset. Accessed: 2024-08-25. Available from: https://pmc.ncbi.nlm.nih.gov/tools/openftlist/.

[14] Lee K, Wei CH, Lu Z. Recent advances of automated methods for searching and extracting genomic variant information from biomedical literature. Briefings in Bioinformatics. 2021;22(3):bbaa142.

[15] Allot A, Peng Y, Wei CH, Lee K, Phan L, Lu Z. LitVar: a semantic search engine for linking genomic variant data in PubMed and PMC. Nucleic Acids Research. 2018;46(Web Server issue):W530–W536.

[16] Allot A, Wei CH, Phan L, Hefferon T, Landrum M, Rehm HL, et al. Tracking genetic variants in the biomedical literature using LitVar 2.0. Nature Genetics. 2023;55(6):901–903.

[17] Gargano MA, Matentzoglu N, Coleman B, Addo-Lartey EB, Anagnostopoulos AV, Anderton J, et al. The Human Phenotype Ontology in 2024: phenotypes around the world. Nucleic Acids Res. 2024;52(D1):D1333–D1346.

[18] Sherry ST, Ward MH, Kholodov M, Baker J, Phan L, Smigielski EM, et al. dbSNP: the NCBI database of genetic variation. Nucleic Acids Research. 2001;29(1):308.

[19] Landrum MJ, Lee JM, Riley GR, Jang W, Rubinstein WS, Church DM, et al. ClinVar: public archive of relationships among sequence variation and human phenotype. Nucleic Acids Research. 2014;42(Database issue):D980–D985.

