## supplementary.pdf for "VUSVista: Enhancing the Curation of Variants of Uncertain Significance"

**4 VUSVista's Evaluation** **10**

**5 List of Abbreviations** **10**

S7 Screenshot from VUSVista of email example of publications update . . 10

#### List of Tables

S1 Accepted user input fields . . . 2

### 1 VUS Submission

| Input Field | Is Required | Accepted Input |
|---|---|---|
| Chromosome number | yes | one number from 1 to 22 or X or Y |
| Variant position on chromosome (locus) | yes | example: 21229389 |
| Reference allele | yes | string made up of: 'G', 'A', 'C', 'T' or '/' |
| Alternate allele | yes | string made up of: 'G', 'A', 'C', 'T' or '/' |
| Zygosity | yes | "heterozygous" or "homozygous" or string in the format: [reference allele]/[alternate allele] (where the alleles can be left empty) |
| Gene symbol | yes | example: "BRCA1" |
| Variant type | yes | examples: "SNV", "insertion", "deletion" |
| Sample identifiers | yes (minimum 1) | example: "sample1" |
| Sample phenotypes | no | examples: "Hypertrophic Cardiomyopathy", "Prolonged Qt Interval" |
| Reference SNP Identification (rsID) | no | example: "rs429358" |
| Human Genome Variation Society (HGVS) nomenclature | no | example: "NM_004168.4:c.1337T¿C" |
| American College of Medical Genetics and Genomics (ACMG) criteria | no | one or more from: PS2, PM3, PM6, PP1, PP4, BS4, BP2, PS3, PP5, BP6, PVS1, PS1, BS3, PM1, BP3, PM2, PM4, PM5, PP2, BP1, PP3, BP4, BA1, BS1, BS2, BP7, PS4, BP5 |
| Publication links | no | example: "https://pubmed.ncbi.nlm.nih.gov/25741868/" |

**Table S1**: **Accepted user input fields.** The variant's position on the chromosome needs to be based on the GRCh37 reference genome. Sample phenotypes need to be from the Human Phenotype Ontology (HPO) to remove the risk of human error on input. The HGVS nomenclature allows scientific members of staff to store the transcript where the variant is found. When comparing recorded variants with newly inputted variants the mandatory fields are compared, excluding the sample identifiers and the zygosity, since they are sample-specific.

#### 1.1 Bulk Upload View

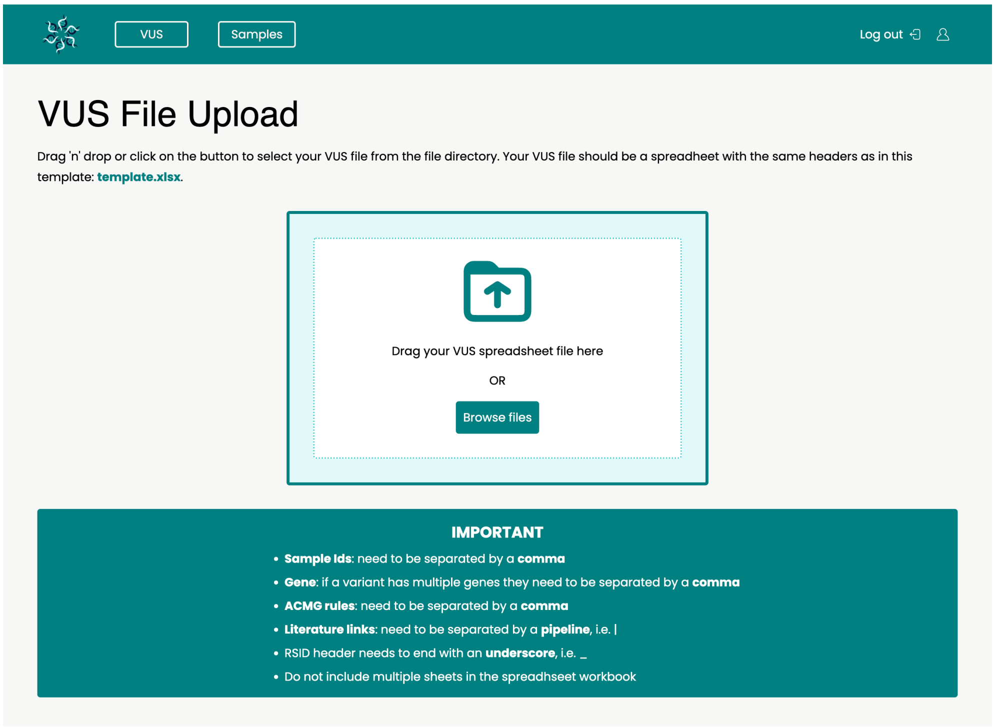

**Fig. S1**: Screenshot from VUSVista of the bulk upload view where users can upload an Excel file containing previously recorded Variants of Uncertain Significance (VUS).

#### 1.2 Variant Processing

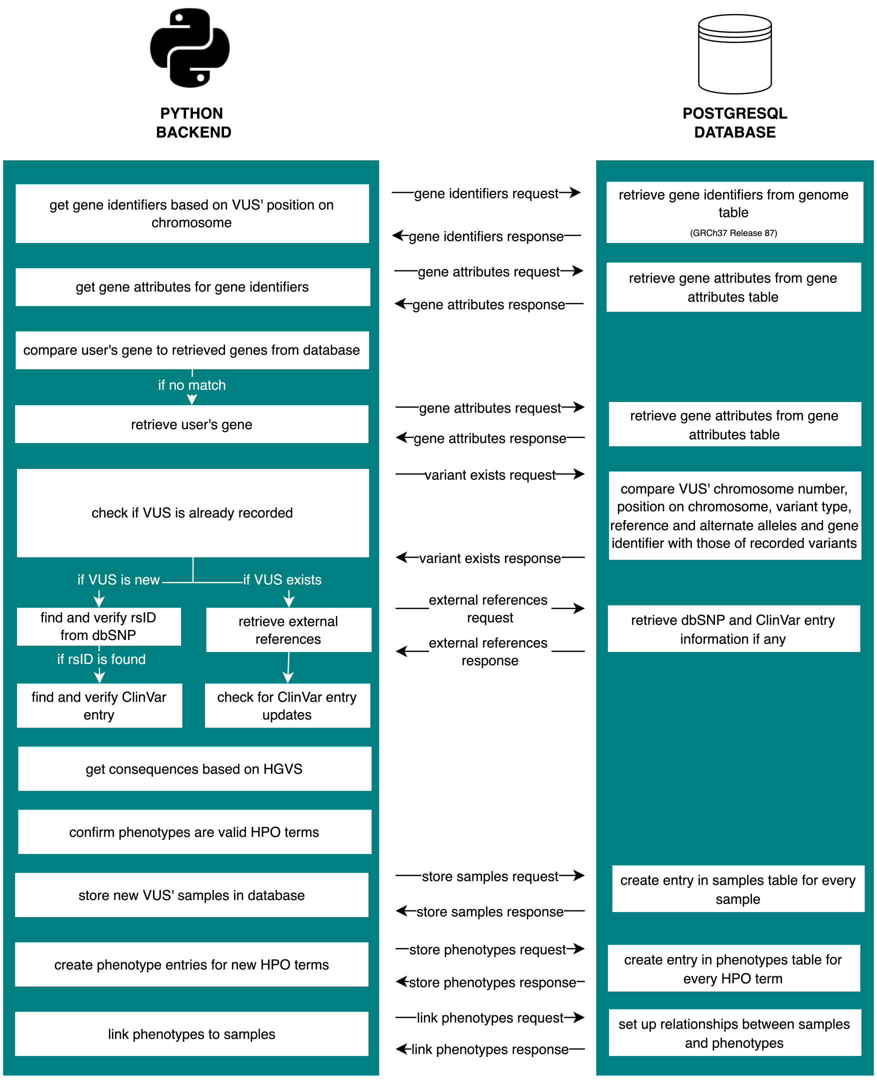

**Fig. S2a:** Retrieval of VUS-related information from external resources. The variant's gene is first retrieved from the database followed by retrieval of external references related to the variant. This is followed by the retrieval of the variant's consequences based on the HGVS nomenclature and the creation and linking of phenotypes and samples.

**Fig. S2**: VUSVista's variant processing

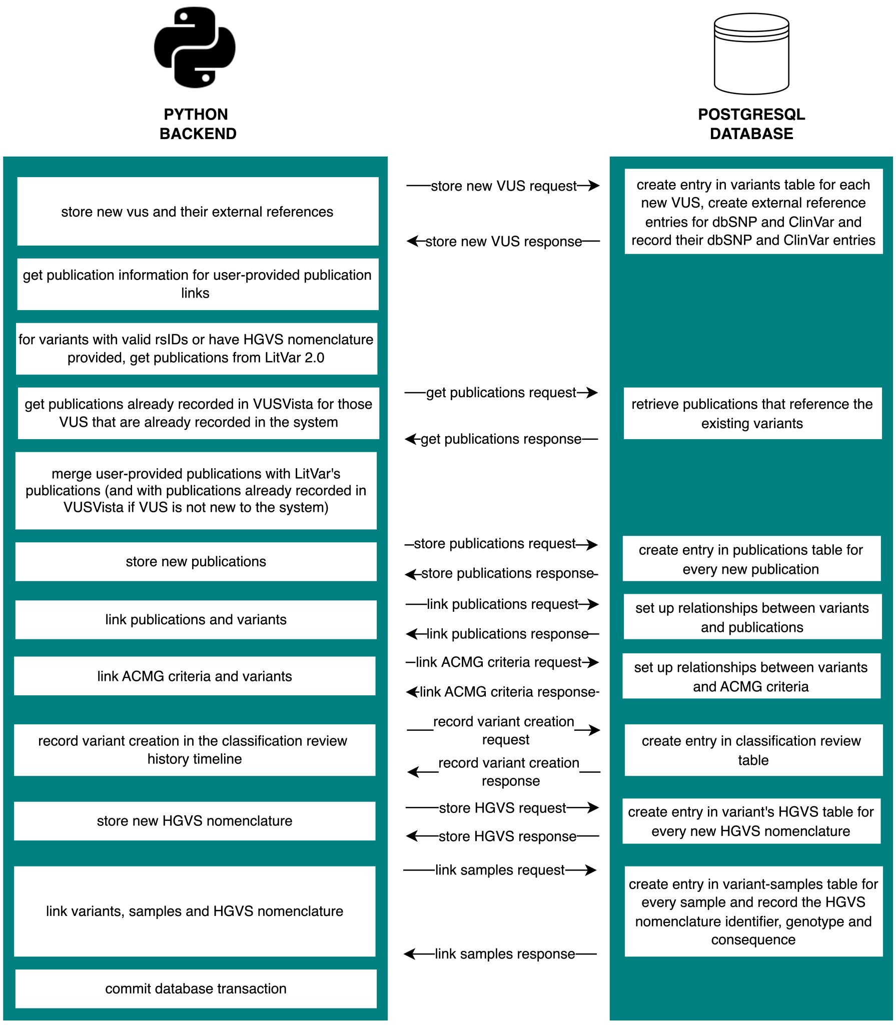

**Fig. S2b**: Storing relevant VUS information in the database. The VUS which are new to the system are recorded together with their external references. Publications are retrieved and for those variants that were already stored in the database, their publications are merged. Publications are linked to variants followed by the ACMG criteria. Finally, the samples and variants relationships are recorded along with their HGVS nomenclatures, genotypes and consequences. The database transaction is then committed.

**Fig. S2**: VUSVista's variant processing

### 2 Variants

#### 2.1 Variant View

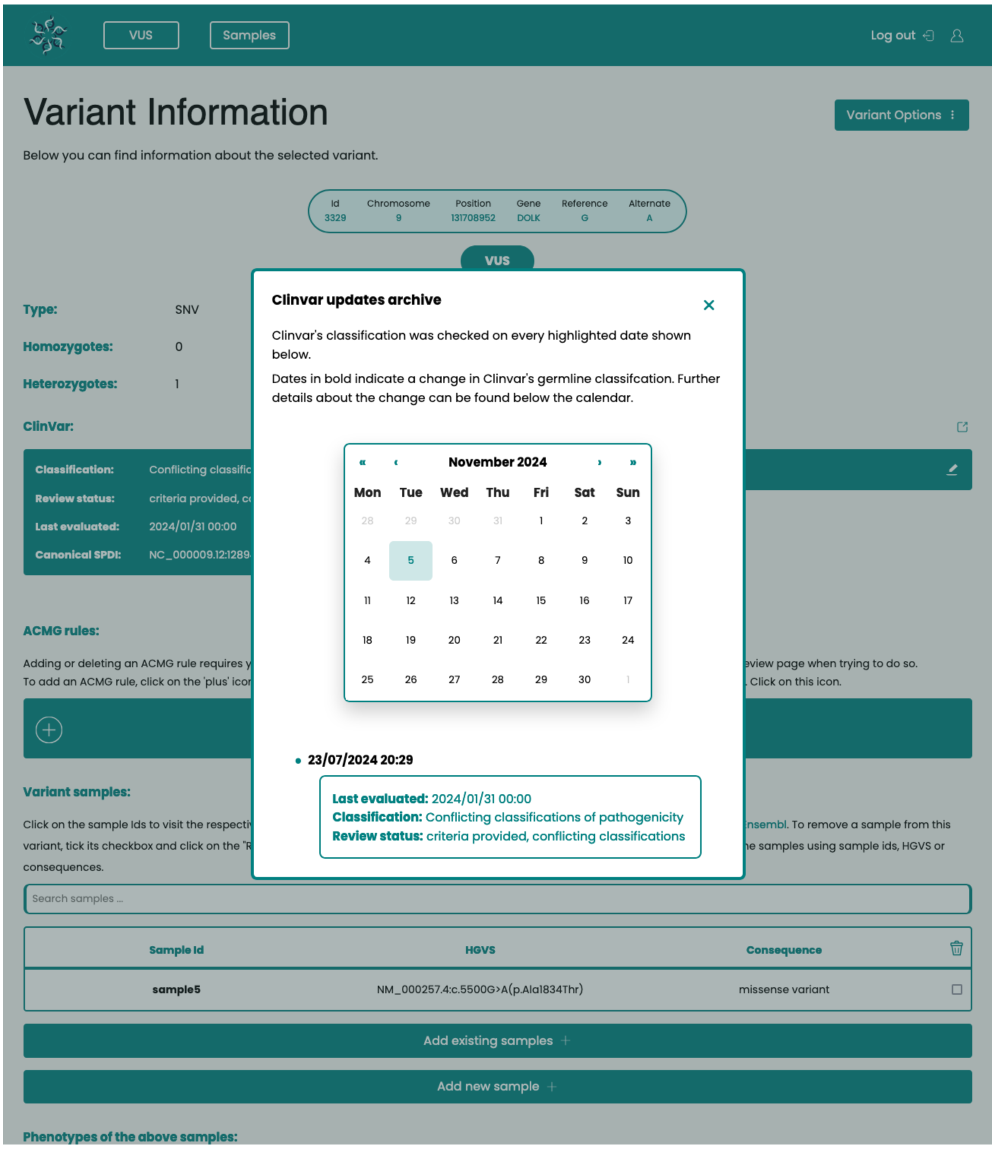

**Fig. S3**: Screenshot from VUSVista of the ClinVar updates archive consisting of calendar marking automatic checks for updates in the variant's ClinVar entry, as well as a timeline showing what changes were made.

### 3 Automated Checks

#### 3.1 ClinVar

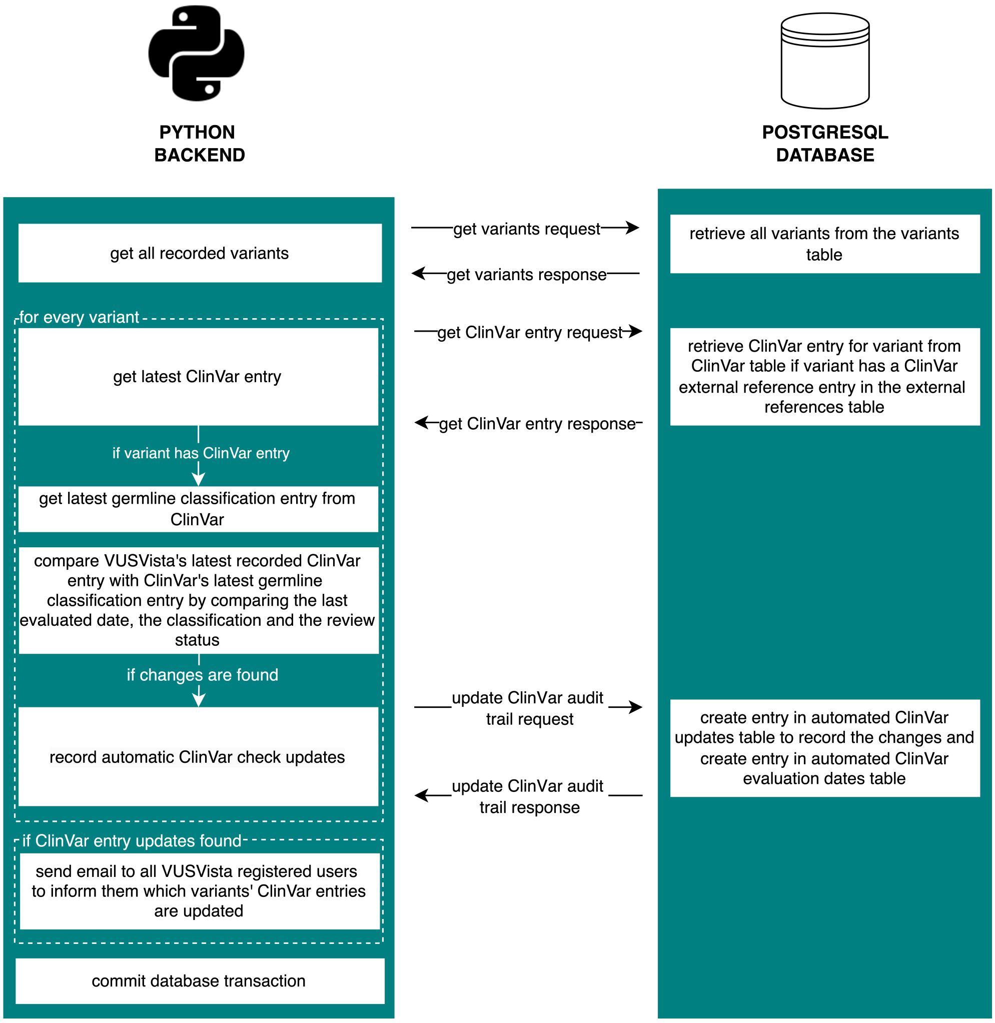

**Fig. S4**: VUSVista's process for automated ClinVar entry update checks. For all recorded variants, VUSVista first retrieves its last recorded ClinVar entry and then retrieves ClinVar's latest germline classification entry. Both entries are compared and if changes are found the variant's ClinVar audit trail is updated, with both the changes found and the date on which the check was executed. If at least one variant's ClinVar entry is updated, an email is sent out to inform users about the changes. Database transaction is finally committed.

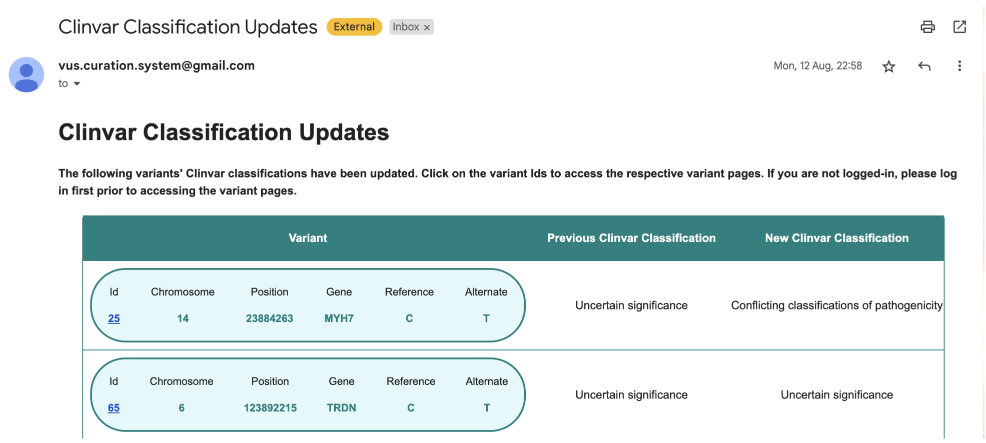

**Fig. S5**: Screenshot from VUSVista of email example of ClinVar update which is sent out if for at least one variant, fields from the ClinVar entry's germline classification, including the classification, the last evaluation date or the review status, are updated.

##### 3.2 Publications

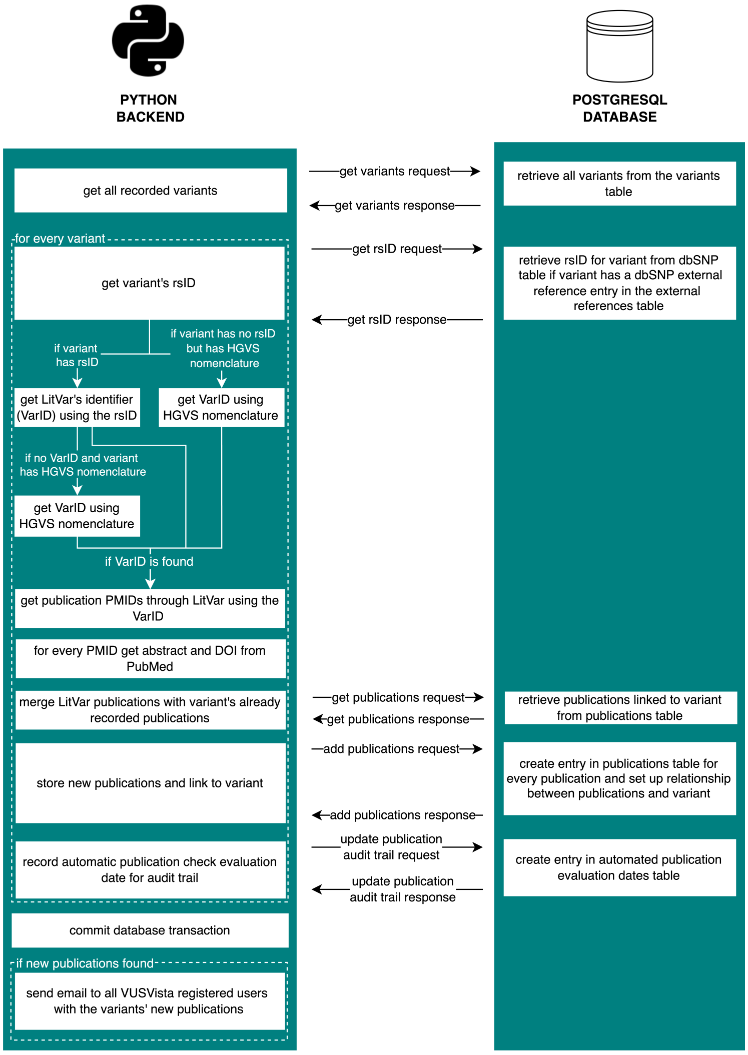

**Fig. S6**: VUSVista's process for automated publication checks. For all recorded variants, VUSVista retrieves LitVar's unique identifier (VarID). PMIDs for publications that reference the variants are first retrieved followed by the retrieval of additional publication information from PubMed. The publication audit trail is then updated. Database transaction is committed and an email is sent out to inform users of newly discovered publications.

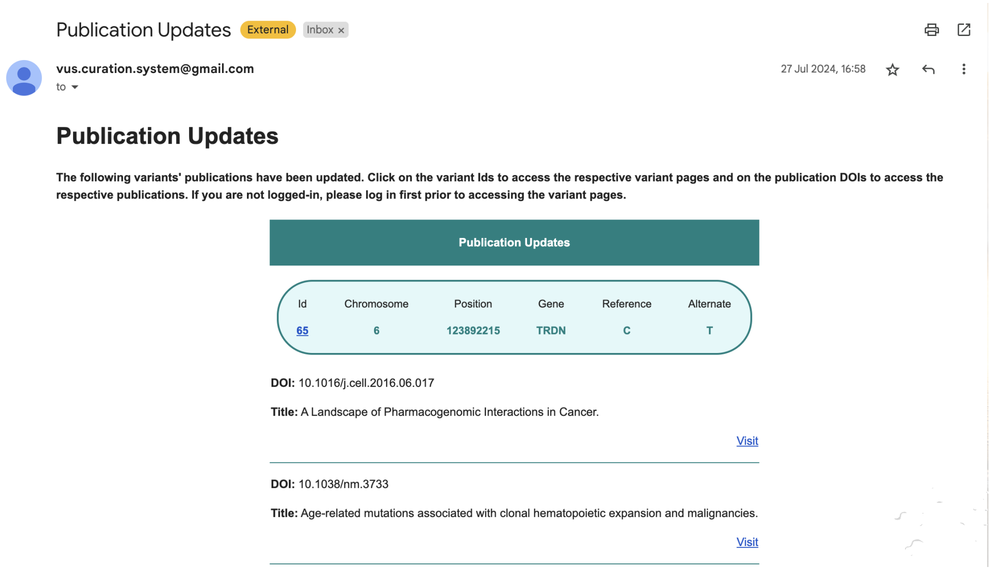

**Fig. S7**: Screenshot from VUSVista of email example of publications update which is sent out if for at least one variant, new publications are found that reference the variant itself, either in the main text or in the publications' supplementary material.

### 4 VUSVista's Evaluation

Focus groups are discussions on a specific topic, led by a researcher, that produce qualitative insights into participants' perspectives and interactions. We recruited participants for individual one-to-one sessions and a focus group. All were medical laboratory scientists or senior pharmacists who encounter VUS in their daily work at a laboratory of molecular pathology and genetics.

During the one-to-one sessions, participants were introduced to VUSVista's features, completed predefined tasks to familiarise themselves with the system, and received the system's user manual (VUSVistaManual.pdf). The focus group was held a few weeks after these sessions, giving participants time to explore VUSVista. Throughout the focus group, we asked the participants about their experience with VUS and their feedback on our system. They offered valuable insights on how effective VUSVista would be if it were to be implemented in their daily work schedule.

### 5 List of Abbreviations

***ACMG*** American College of Medical Genetics and Genomics

***HGVS*** Human Genome Variation Society

***HPO*** Human Phenotype Ontology

***rsID*** Reference SNP Identification

***VUS*** Variants of Uncertain Significance
