## Supplementary material for "VUSVista: Enhancing the Curation of Variants of Uncertain Significance": VUSVistaManual.pdf

#### VUSVista Manual

Access from: [https://vus-vista.vercel.app/](https://vus-vista.vercel.app/)

### Important Notice

**If any errors occur, please send an email with information about what you were trying to do. If possible provide steps on how to replicate the issue. If the issue is related to the file upload operation, kindly attach the file to the email. Ensure there is no patient data in the file.**

### Login Page

Enter your credentials and click on the log in button. Once you login, you will find yourself in the Homepage.

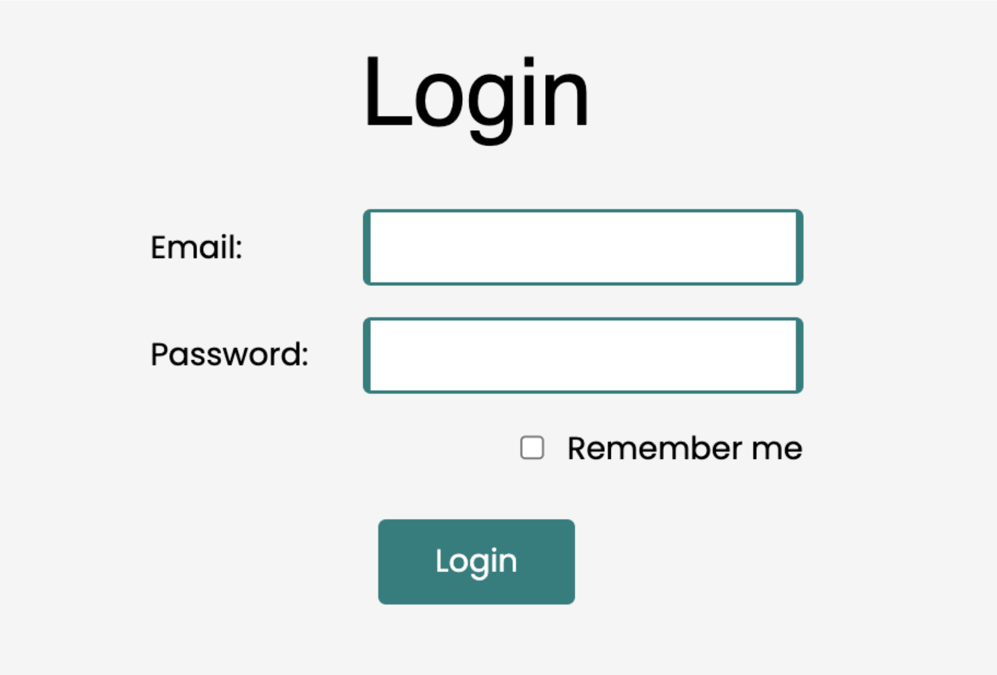

### Homepage

VUSVista

The software automatically checks if there are any changes in the stored variants' Clinvar classifications. This is the last date that the software checked for updates.

**Clinvar last auto update:** 16/07/2024 18:28

**Publications last auto update:** 16/07/2024 18:28

The software automatically checks for any new publications related to the stored variants. This is the date when this check was last carried out.

Most recently uploaded variants

#### Latest uploaded variants

| Variant Id | Chromosome | Position | Gene | Reference | Alternate | RSID |
|---|---|---|---|---|---|---|
| 2740 | 10 | 88439188 | LDB3 | A | G | rs935307586 |
| 2741 | 2 | 21229389 | APOB | T | G | rs769074061 |
| 2742 | 10 | 18439792 | CACNB2 | TT | | |
| 2743 | 5 | 236619 | SDHA | T | C | rs201741295 |
| 2744 | 5 | 236628 | SDHA | C | T | rs201139275 |

### Header

Navigate through the software by accessing the different available pages from the Header.

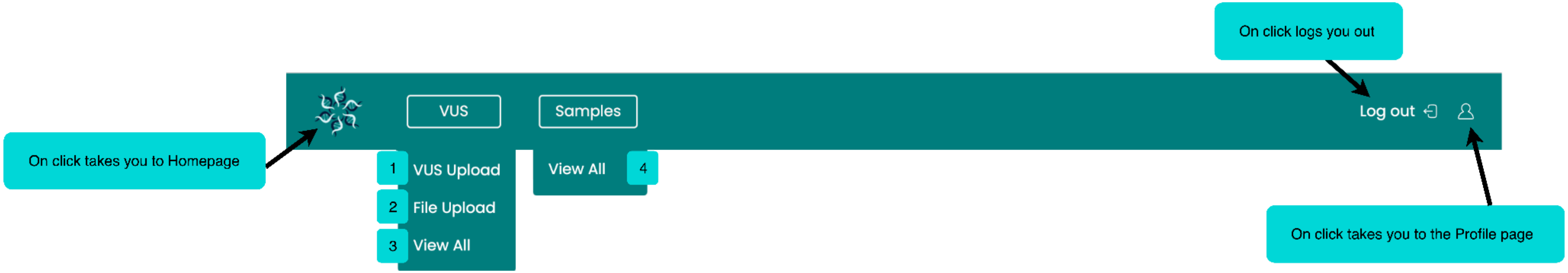

1. Click to visit the VUS Upload page. Here you can manually add a new variant.
2. Click to visit the File Upload page. Here you can upload a spreadsheet with new variant records (bulk upload).
3. Click to visit the View All Variants page. Here you can view all of the variants uploaded by all the users.
4. Click to visit theView All Samples page. Here you can view all of the samples that have any of the uploaded variants.

You can find a section about each of the mentioned pages later in the manual.

### View All Variants Page

This page contains a table with all the variants inputted by all the users making use of this software.

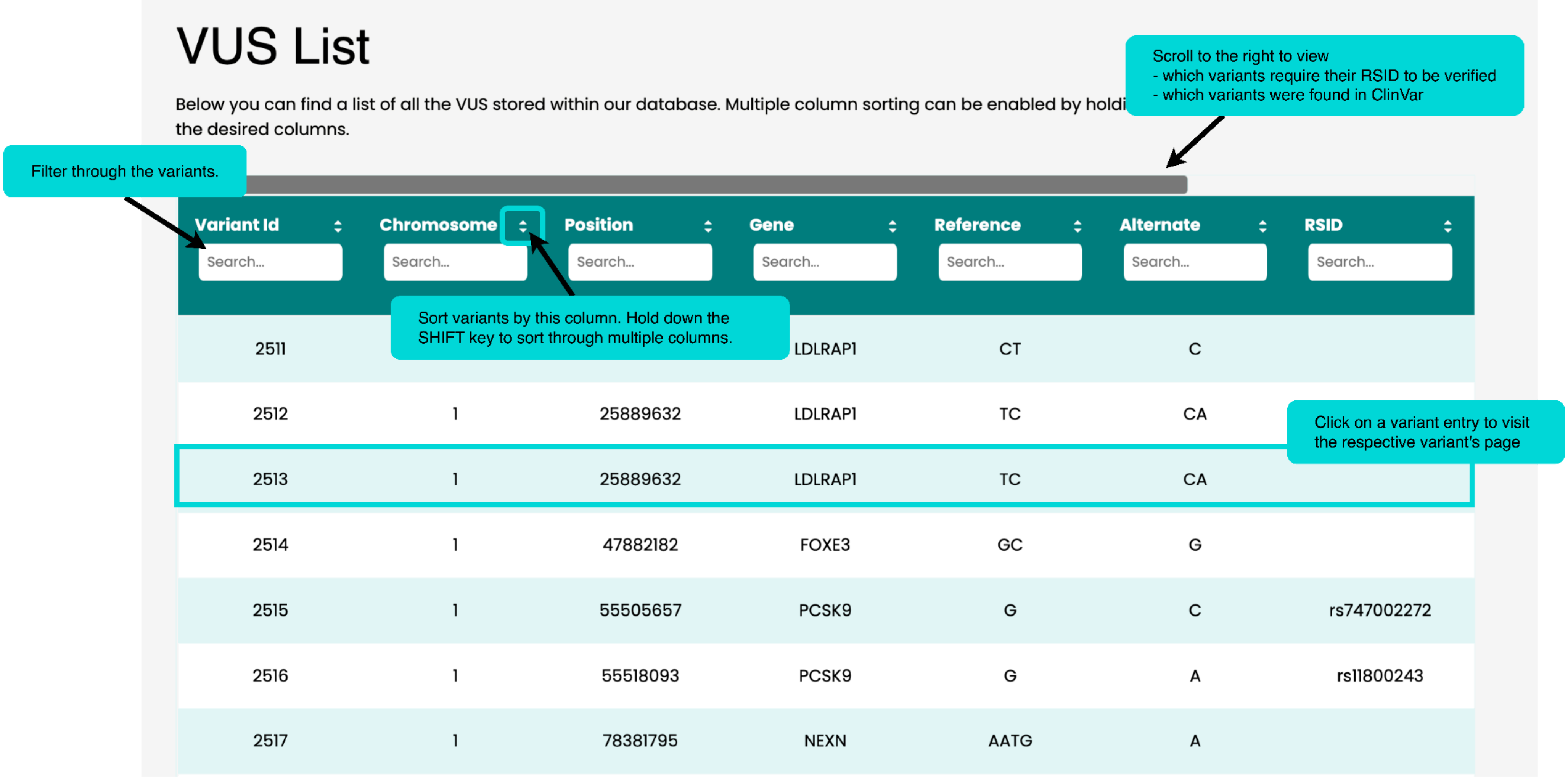

### Variant Page - Part 1

This page is accessible by clicking on a variant entry in the View All Variants page.

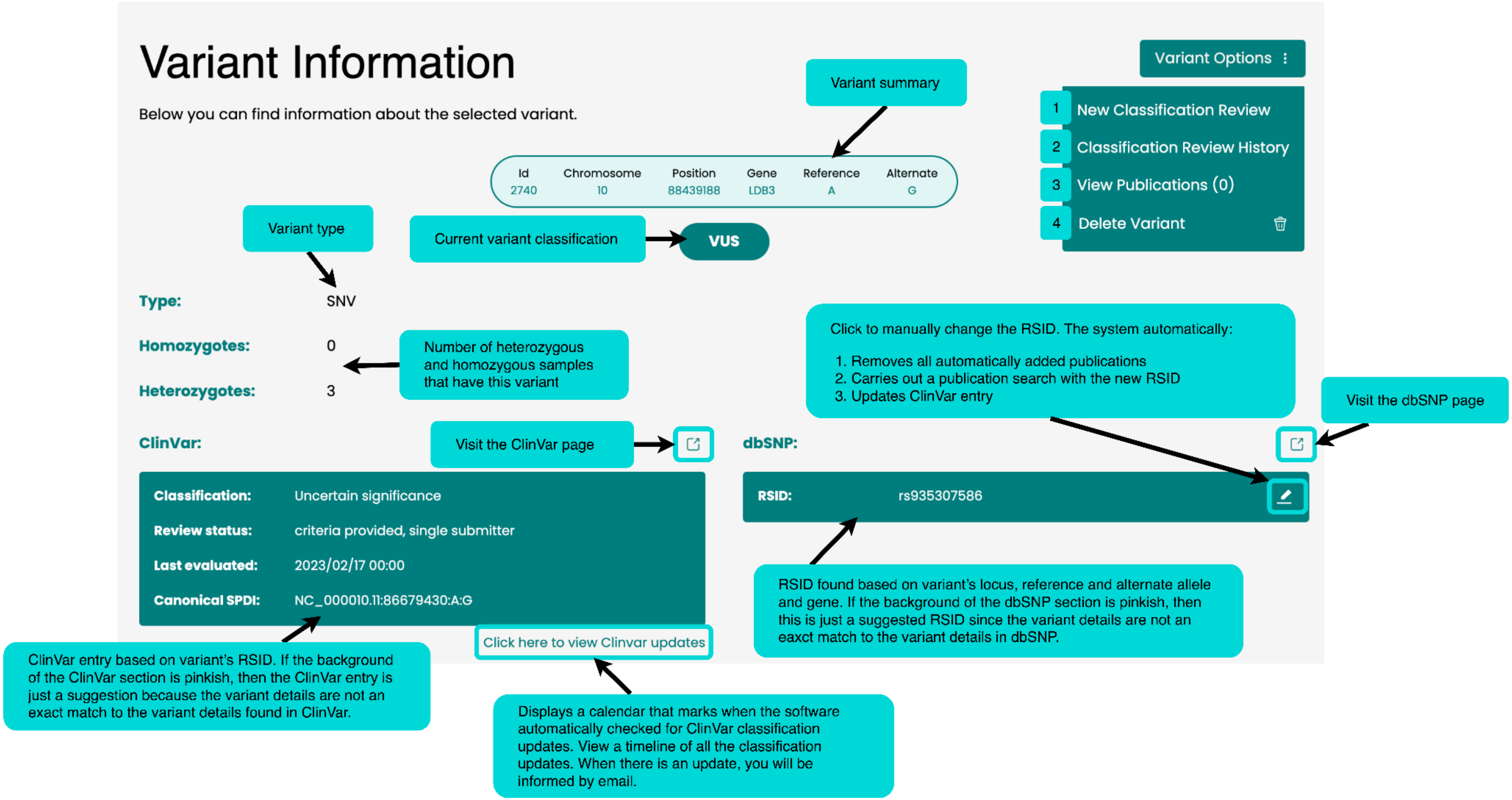

1. Click to visit the New Classification Review page. Here you can add a new classification review either (a.) because you are updating the variant's classification or (b.) because you found relevant information about the variant that you would like to keep record of.
2. Click to visit the Classification Review History page. Here you can view a timeline showing when classification reviews were created.
3. Click to visit the variant's Publication page.
4. Click to delete the variant (*Note:* samples that only have this variant will also be deleted).

You can find a section about each of the mentioned pages later in the manual.

### Variant Page - Part 2

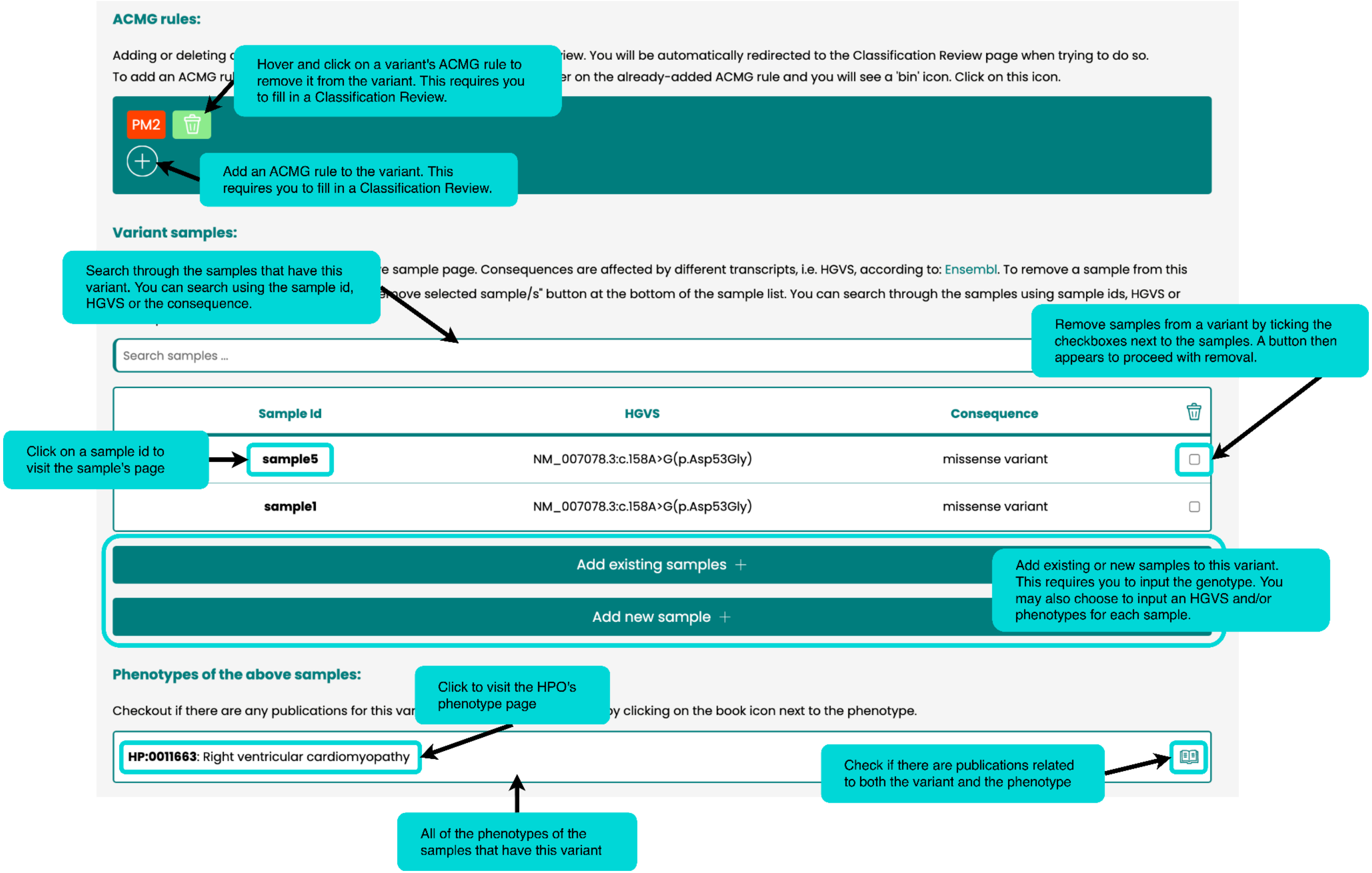

### New Classification Review Page

Create a new classification review for a variant either (a.) because you are updating the variant's classification or (b.) because you found relevant information about the variant that you would like to keep record of.

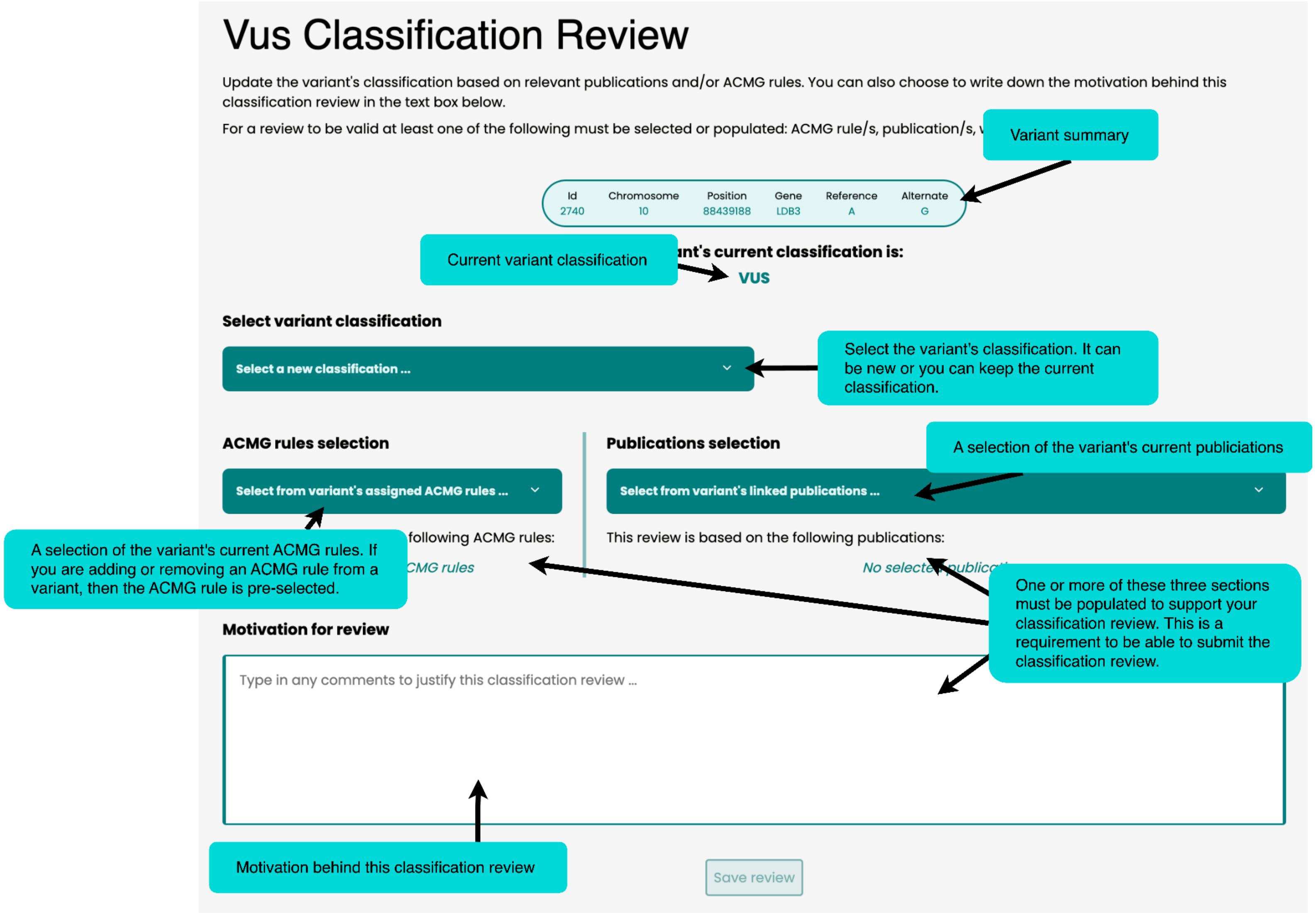

### Classification Review History Page

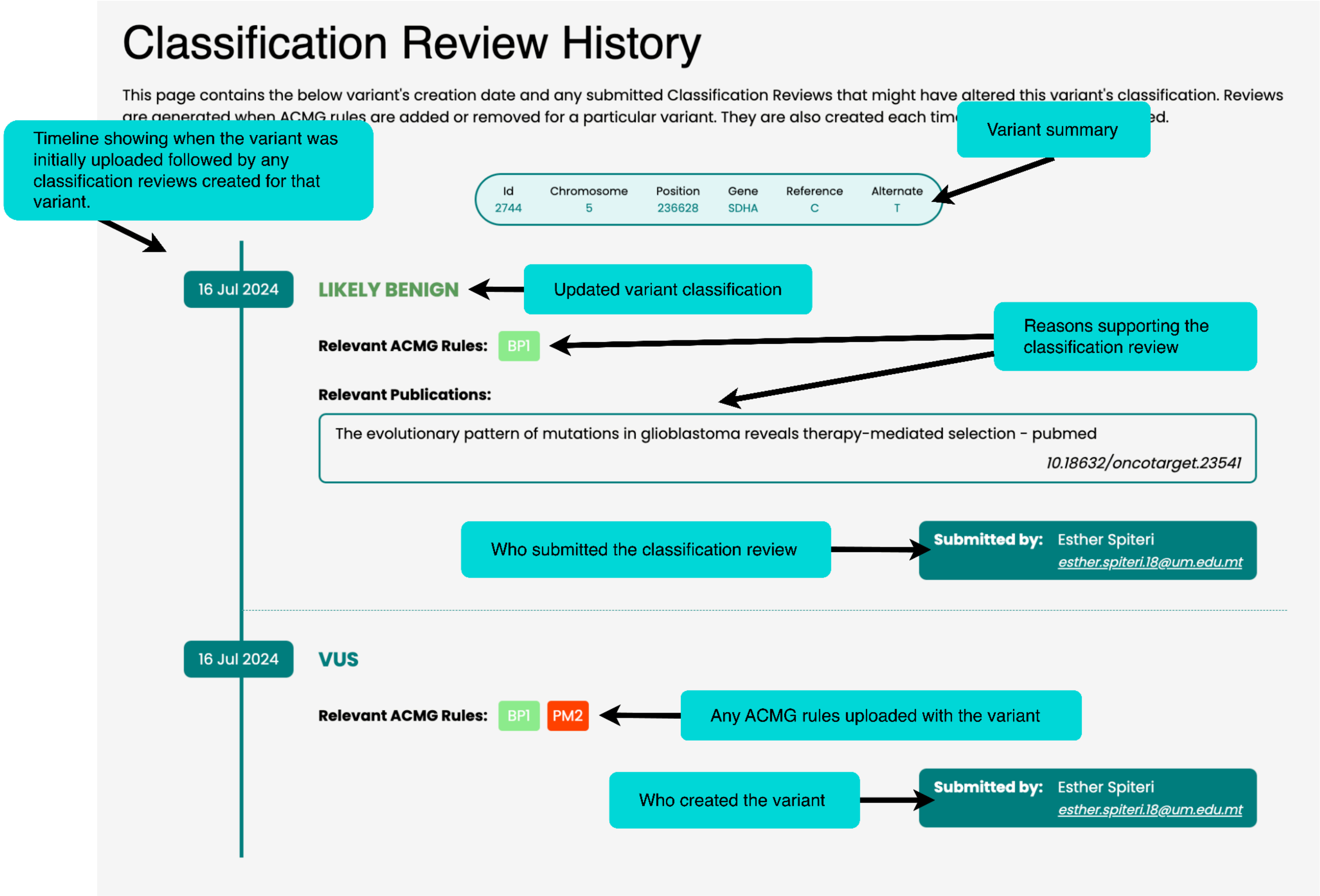

### Publications Page

In this page you will find the variant's publications.

These can be automatically found my the system based on the variant's RSID or one of its HGVS. It is important to note that the software standarizes the variant's nomenclature to search for publications. Therefore, you may need to look up the variant using different aliases in the found publications.

A variant's publications can also be added manually by the user.

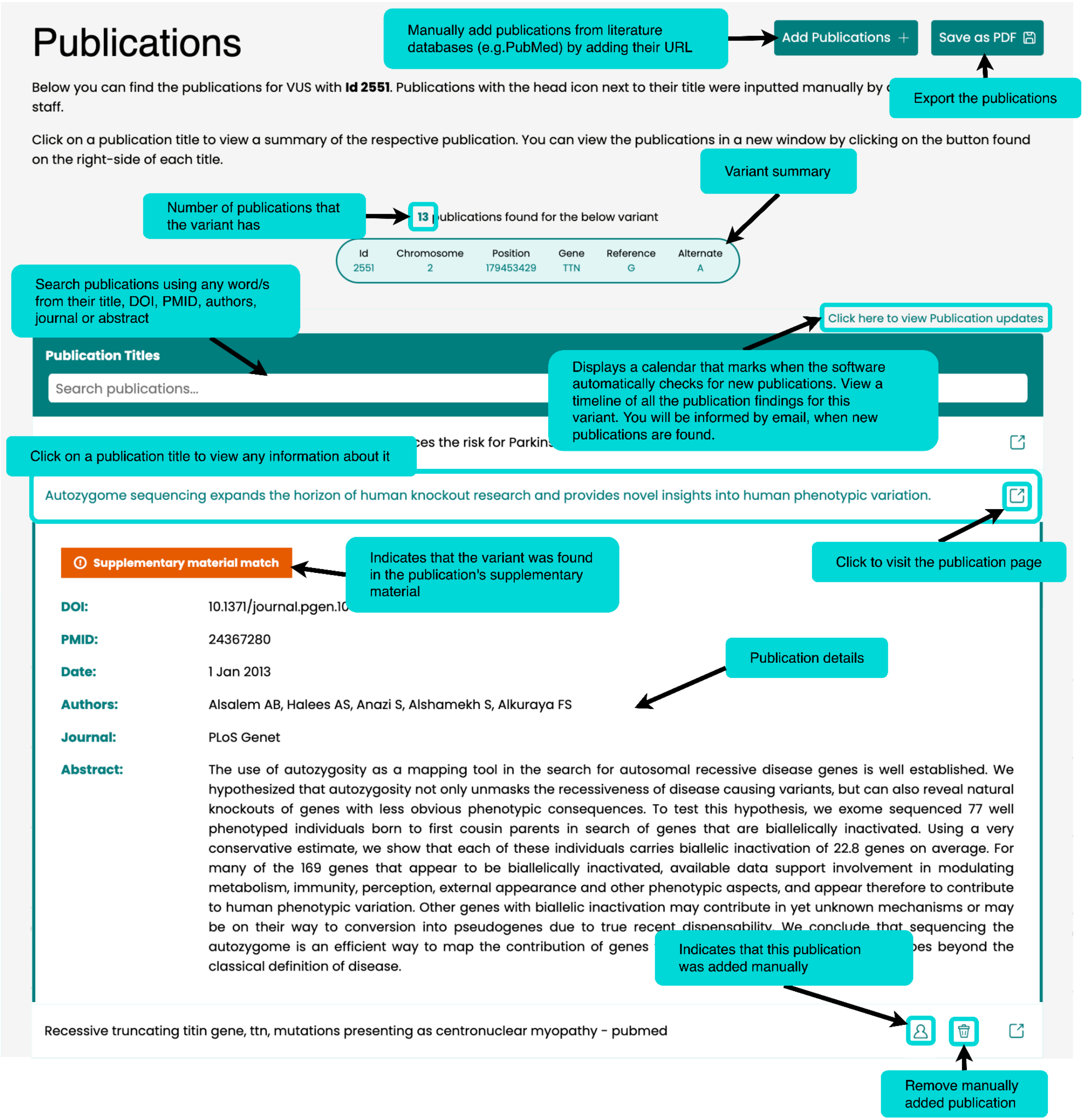

### VUS Upload Page

Manually upload a VUS by filling in the form. It is important to provide information based on the GRCh37 (hg19) build since the software only supports this genome. Variants which had been uploaded in the past do not get overwritten by the newly uploaded data. For existing variants, new samples are added to the variant together with the respective HGVS.

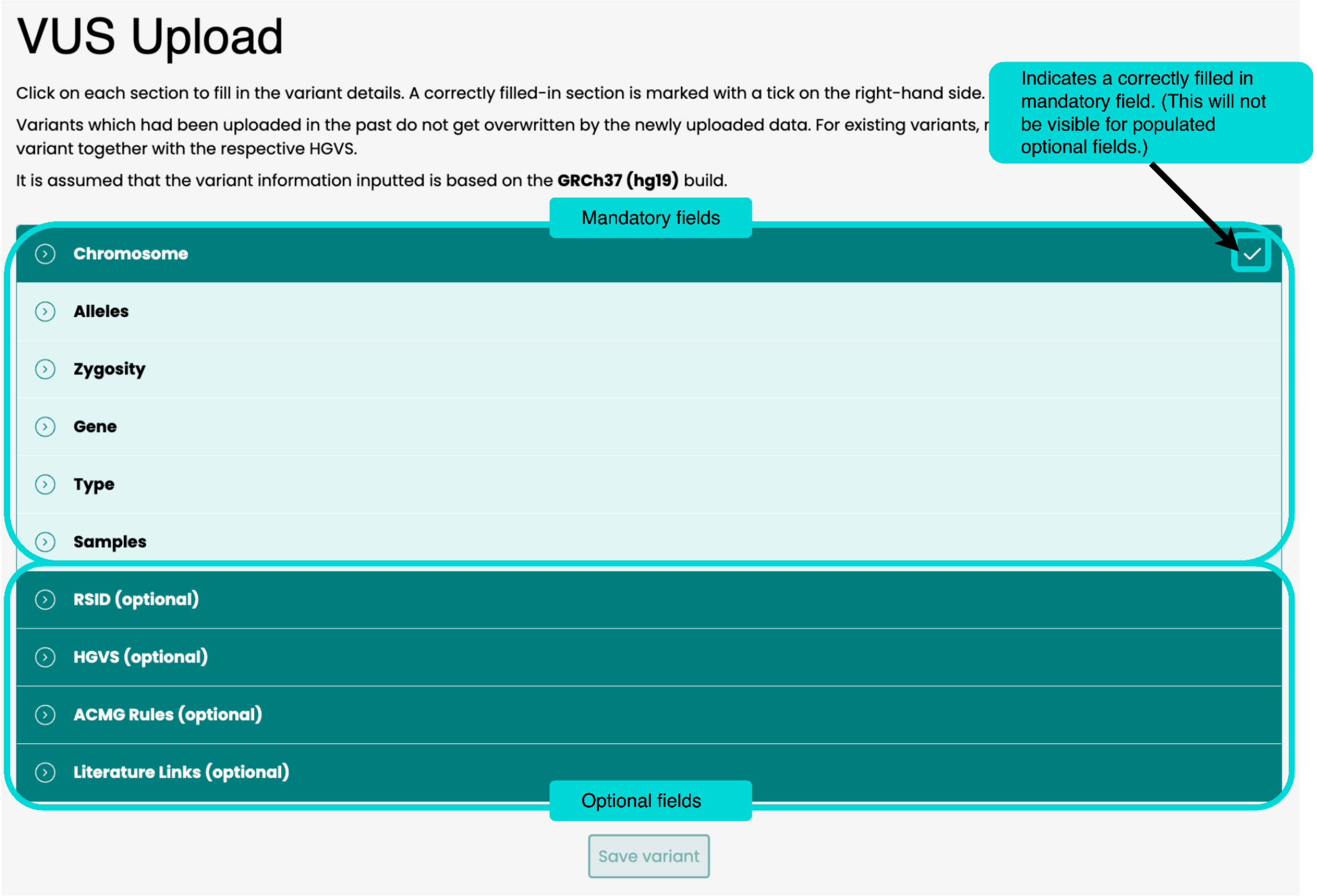

### File Upload Page

In this page you can do a bulk upload of variants in a spreadsheet. A template is provided for download that indicates the spreadsheet's required format. It is populated with example inputs to help guide you with populating your own file.

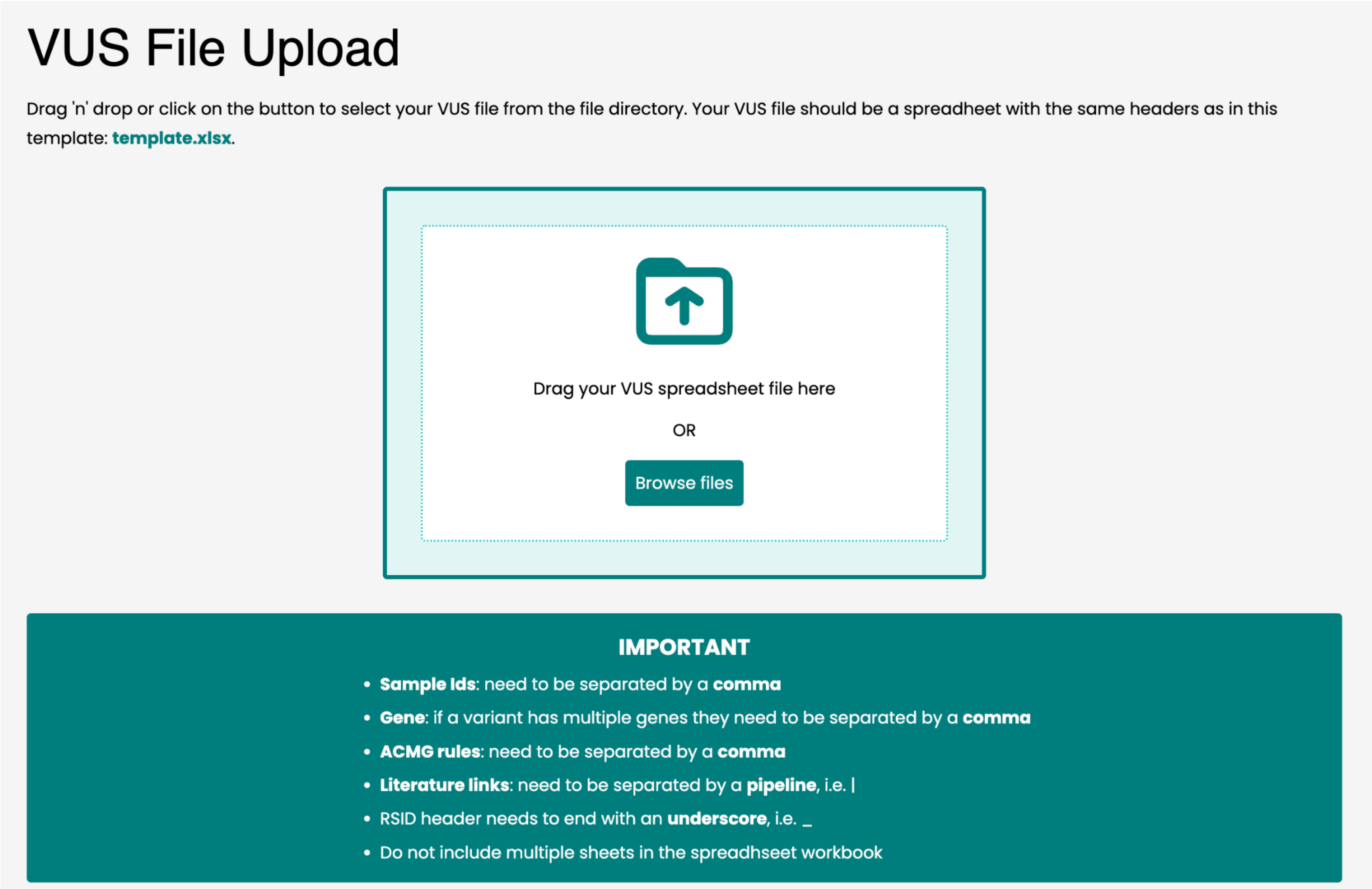

The following checks need to carried out prior to uploading the file:

- A workbook must contain only a single spreadsheet
- Headers must be in the first row
- Headers must be created even if a particular column has no data
- Headers need to match exactly to the following:
    - Sample Ids
    - Locus
    - Gene
    - Type
    - Genotype
    - Reference
    - Alt
    - HGVS
    - RSID_
    - Classification
    - Sample Phenotypes
    - ACMG Rules
    - Literature Links

The following rules apply:

- Sample Ids: separated by comma
- Gene: separated by comma

- ACMG rules: separated by comma
- Literature links: separated by pipeline (|)
- Genotype: can be either in the format [refAllele]/[altallele] or heterozygous or homozygous

Please note:

- If a VUS entry has multiple genes you will be prompted to choose one of them.
- If a VUS entry has genes that are not found in the system database you will be prompted to input a valid gene alias.
- Once you upload the file for processing, you can continue making use of the software. A banner will then appear to inform you about the file’s upload status.

### View All Samples Page

This page displays all of the samples that have at least one of the variant's uploaded to this software.

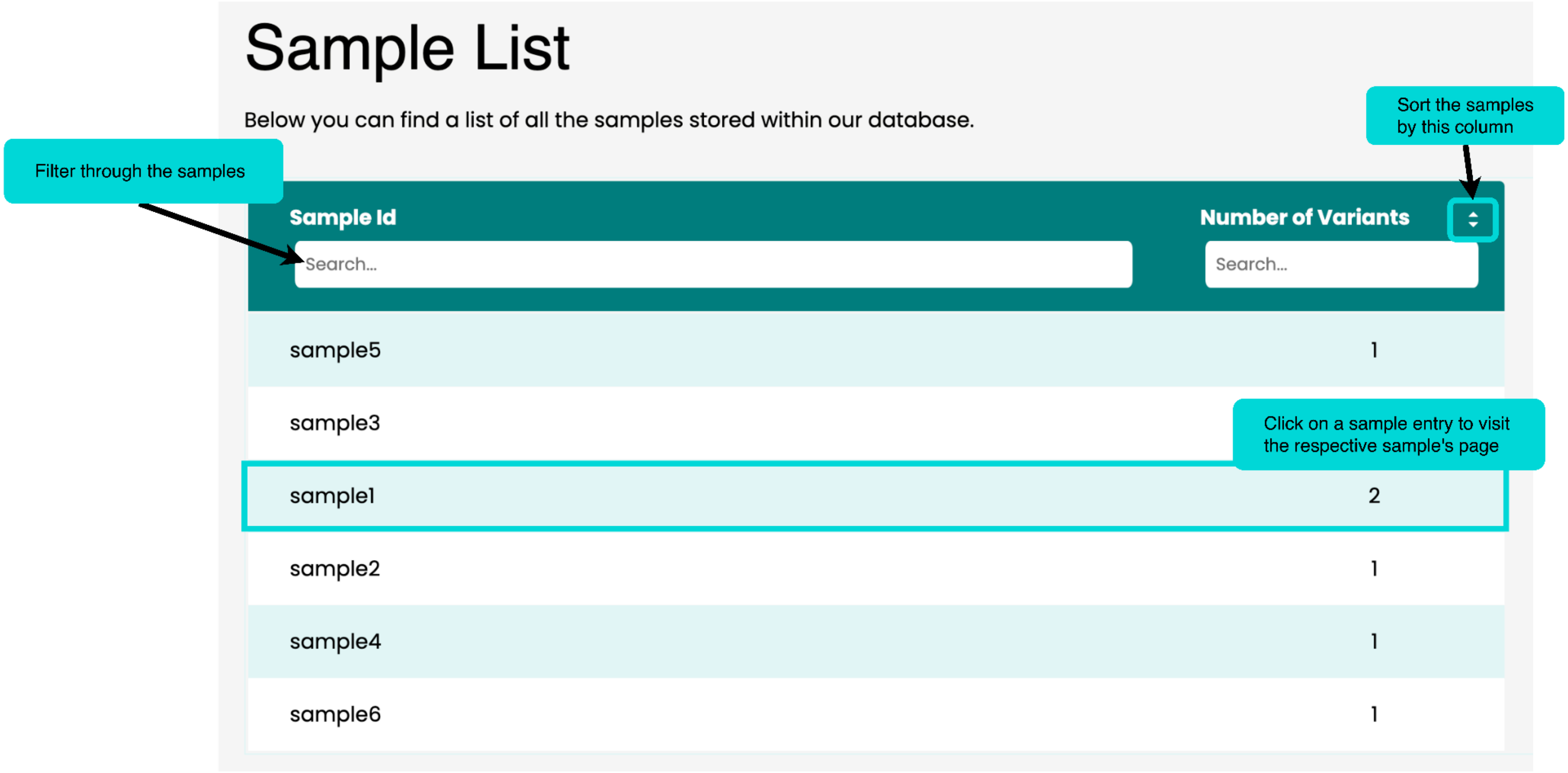

### Sample Page

This page is accessible by clicking on a sample entry in the View All Samples page.

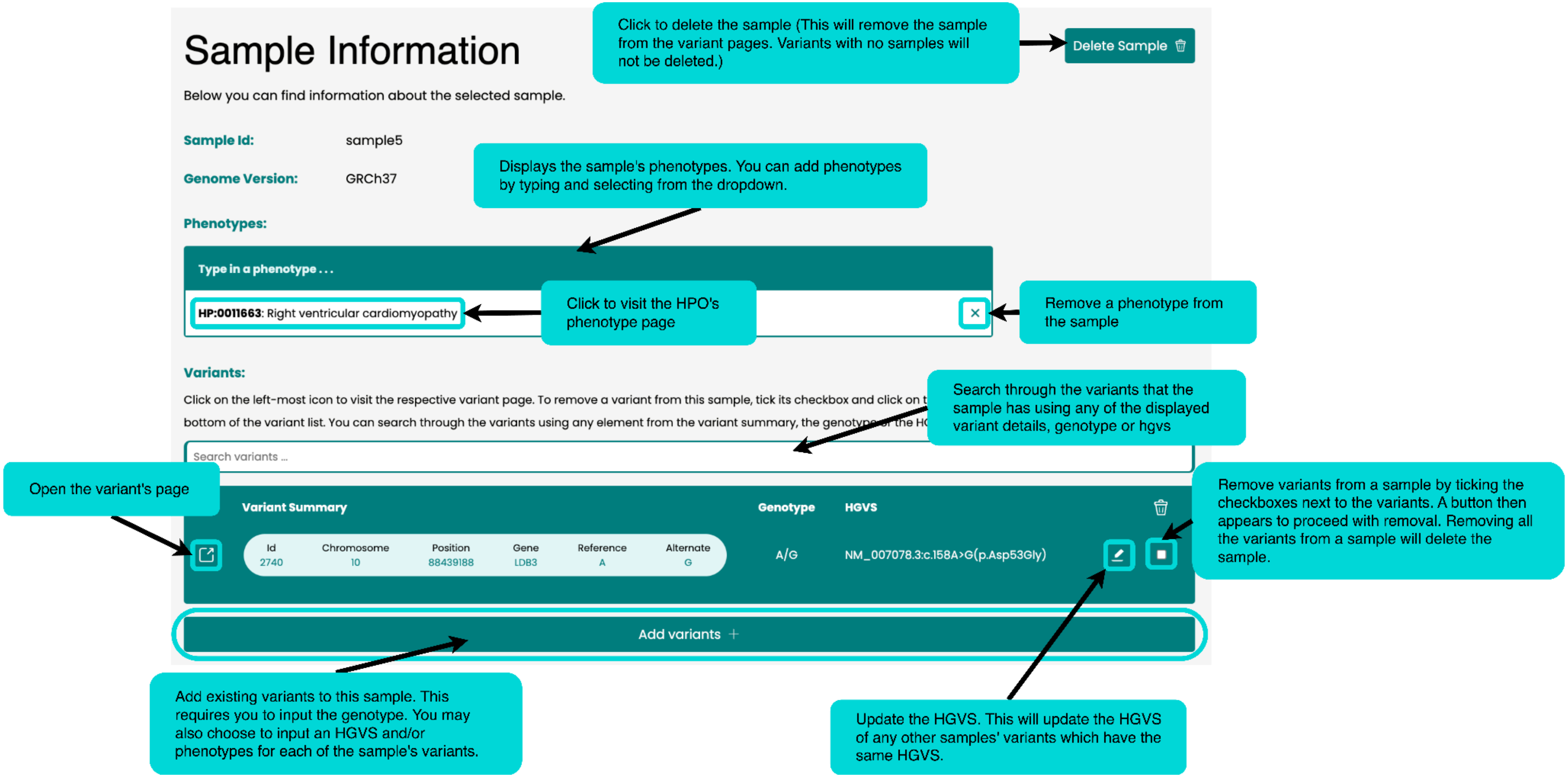

#### Profile Page

Displays your profile's information.

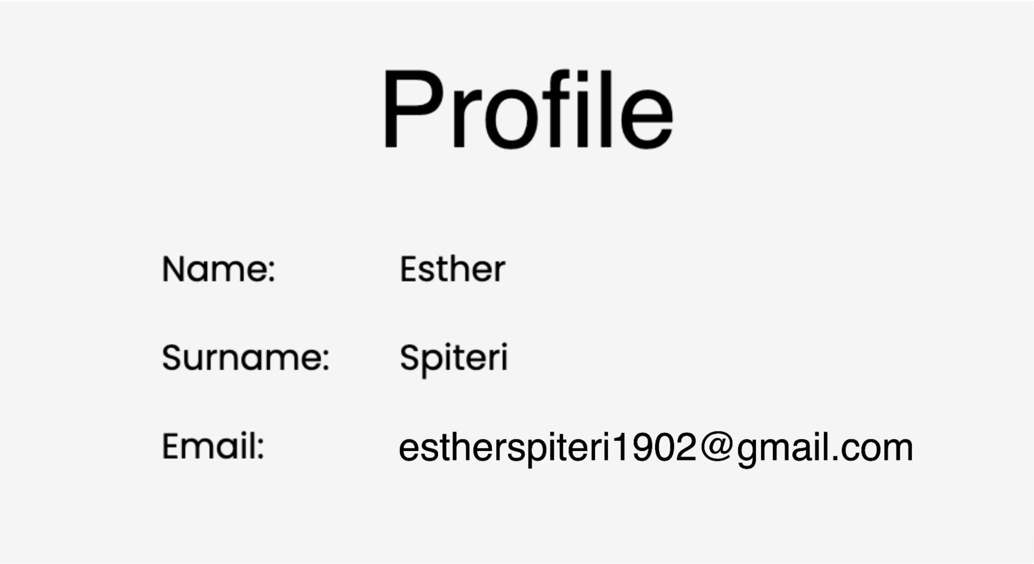
